## Supplemental Files for "Limited within-host diversity and tight transmission bottlenecks limit SARS-CoV-2 evolution in acutely infected individuals"

### Supplementary Materials

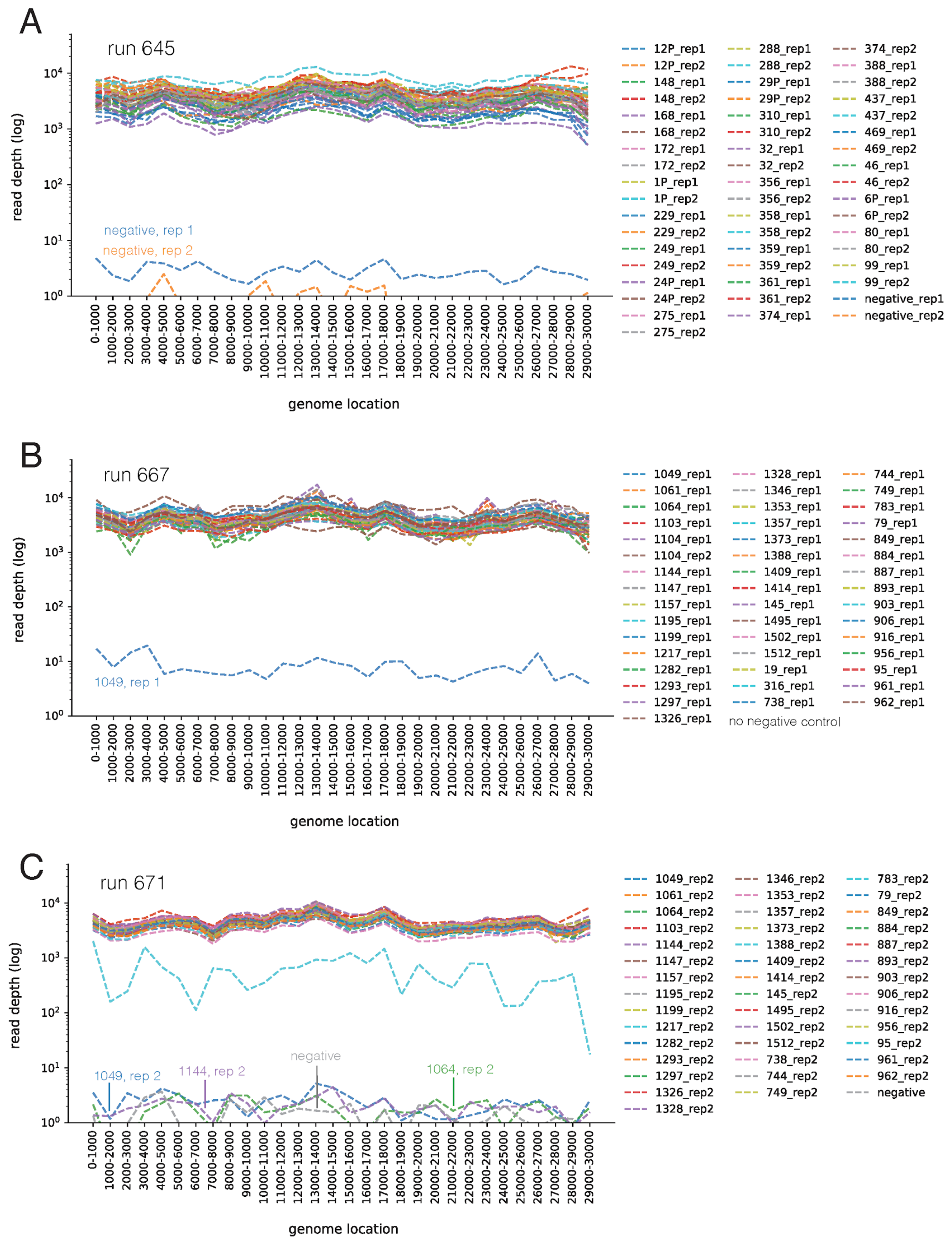


**Supplemental Figure 1: Read depth**

Read depth by genome location in 1,000-bp bins for MiSeq runs **a.** 627, **b**. 628, **c.** 643, and **d.** 644. Water controls and low-coverage samples are labeled. Samples included in each run are labeled according to the color to the right of each plot.

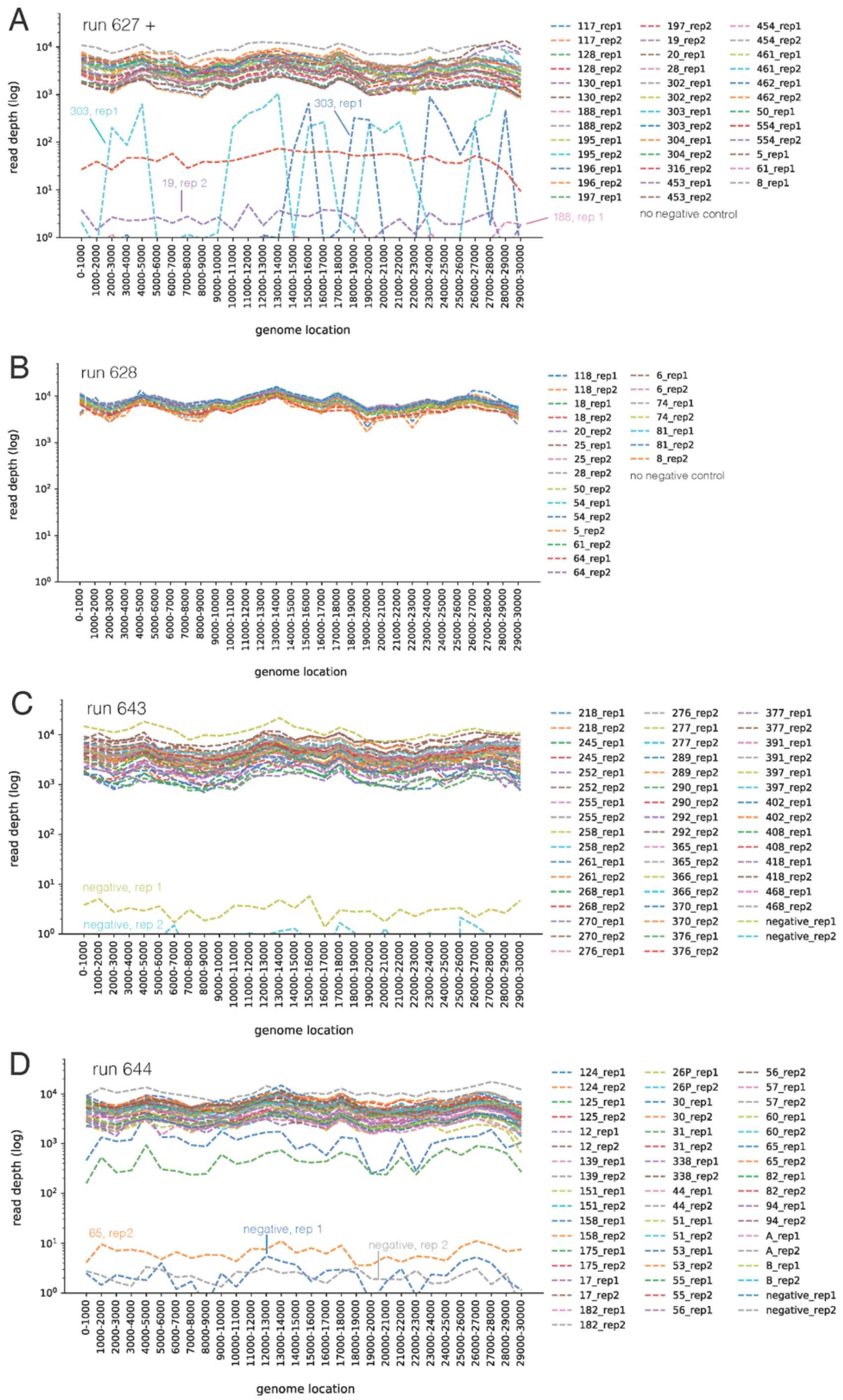


**Supplemental Figure 2: Read depth**

Read depth by genome location in 1,000-bp bins for MiSeq runs **a.** 645, **b**. 667, and **c.** 671. Water controls and low-coverage samples are labeled within each plot. Samples included in each run are labeled according to the color to the right of each plot.

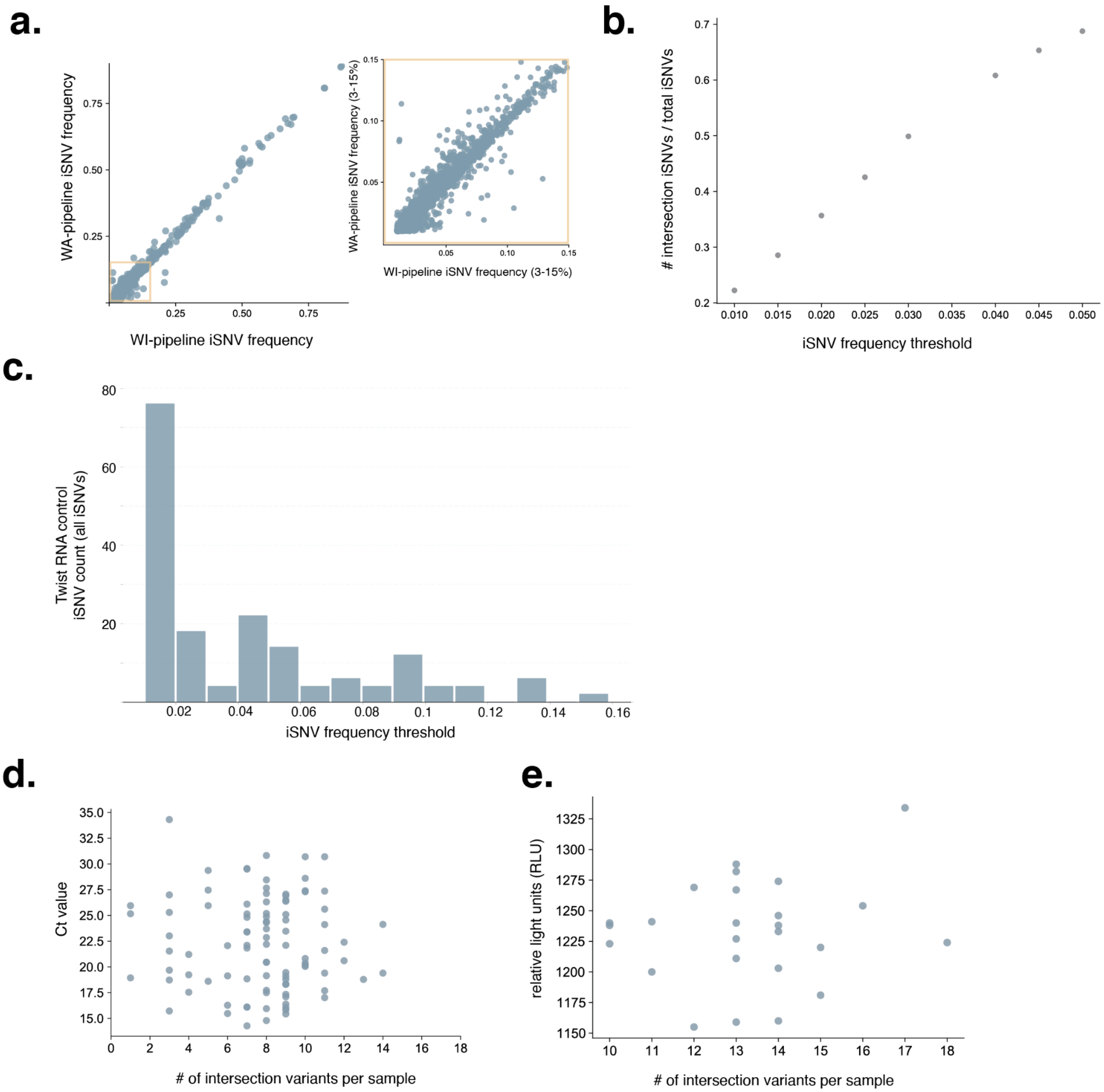


**Supplemental Figure 3: Additional iSNV quality control information**

Subplot **a**. shows variant frequencies generated using the Wisconsin bioinformatic pipelines are shown on the x-axis and frequencies generated using the Washington bioinformatic pipeline are shown on the y-axis. The yellow box highlights low-frequency variants (3-15%), which is expanded out to the right. **b**. Proportion of intersection iSNVs relative to the total number of iSNVs increases as variant frequency threshold increases. **c**. The total number of iSNVs detected across both Twist RNA control replicates compared to the iSNV frequency threshold. The majority of iSNVs detected in these clonal samples occur

<3% frequency. Note that the iSNVs reported in **Supplemental Table 1** are intersection iSNVs only. The identities of all iSNVs detected ≥1% frequency in the Twist RNA control can be found in the GitHub accompanything this manuscript. **d.** The number of intersection variants is compared to the Ct value for all samples where a Ct value was available. Out of 133 total samples, Ct values were available for 94. **e.** The number of intersection variants is compared to the RLU (relative light unit) value for all samples where a RLU value was available.

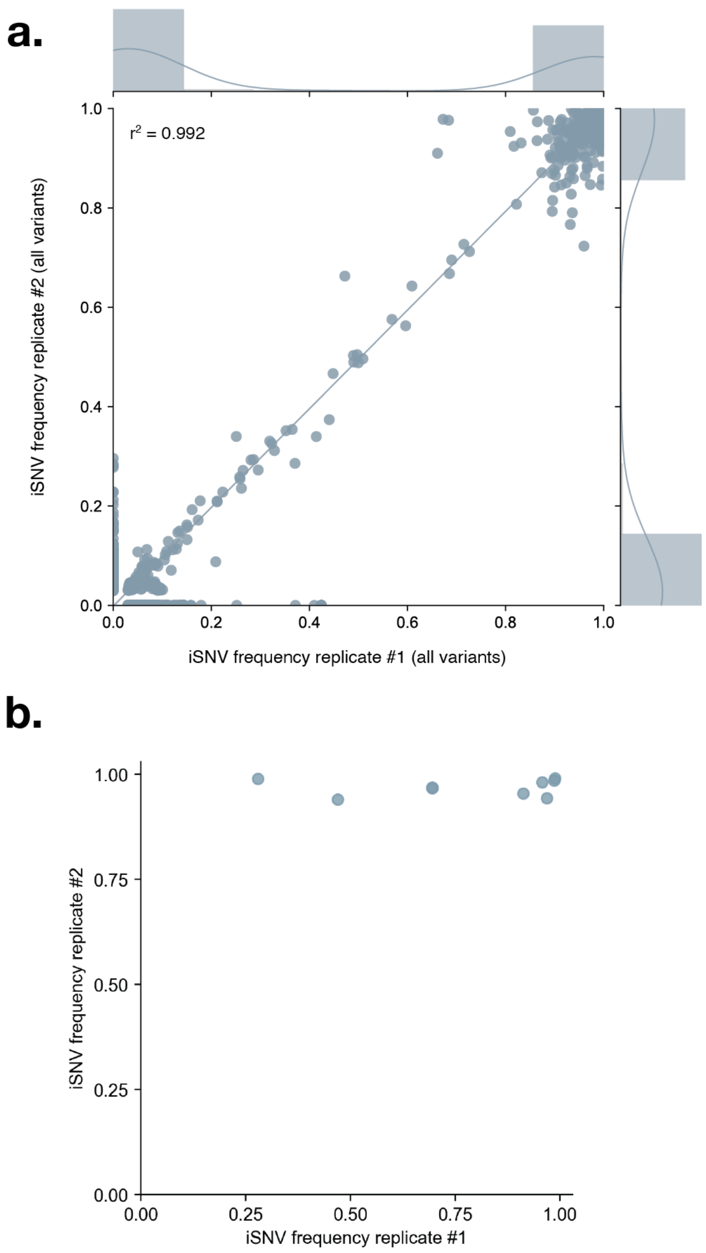


**Supplemental Figure 4: iSNVs in technical replicates across all samples. a.** Variant frequencies in replicate 1 are shown on the x-axis and frequencies in replicate 2 are shown on y-axis. This plot includes all variants found in both replicates and not just the intersection variants as shown **Figure 1a**. **b.** Example of one sample with very poor overlap between technical replicates; this sample (sample 1104) was excluded from the experimental dataset.

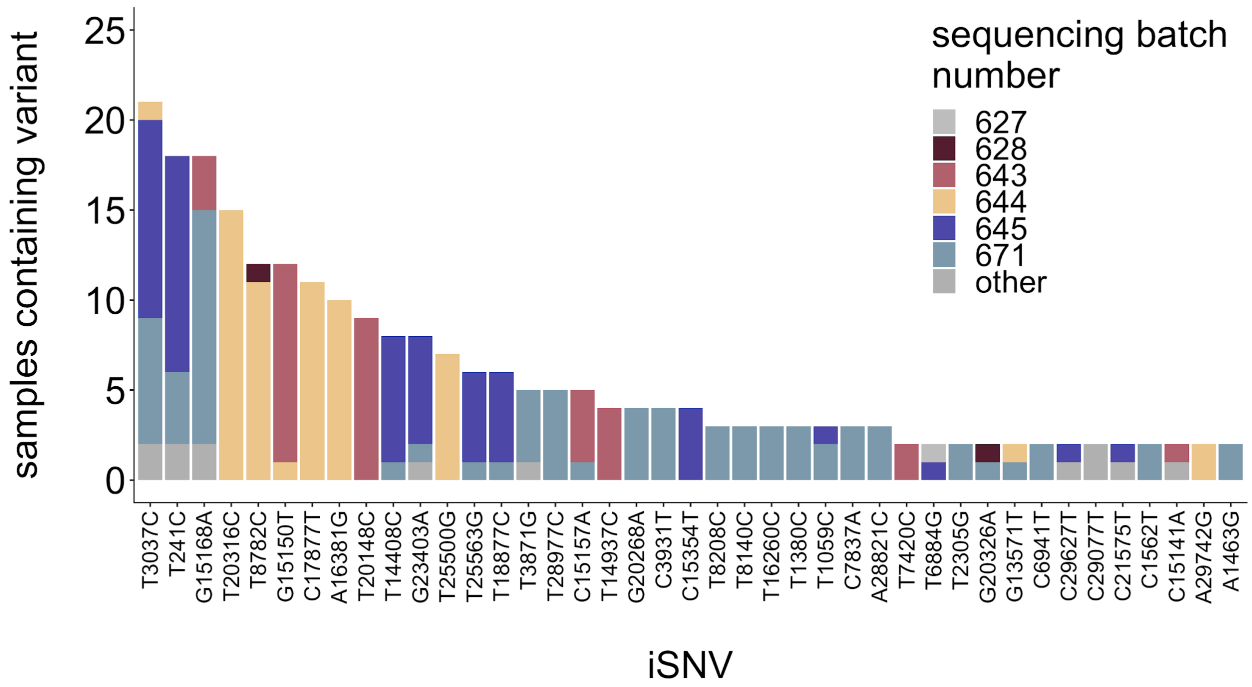


**Supplemental Figure 5: iSNVs do not cluster by sequencing run.** iSNVs detected in at least 2 samples are shown on the x-axis and are plotted against the number of times they are detected in our dataset. Each iSNV bar is colored according to the number of times it was detected within each sequencing batch.

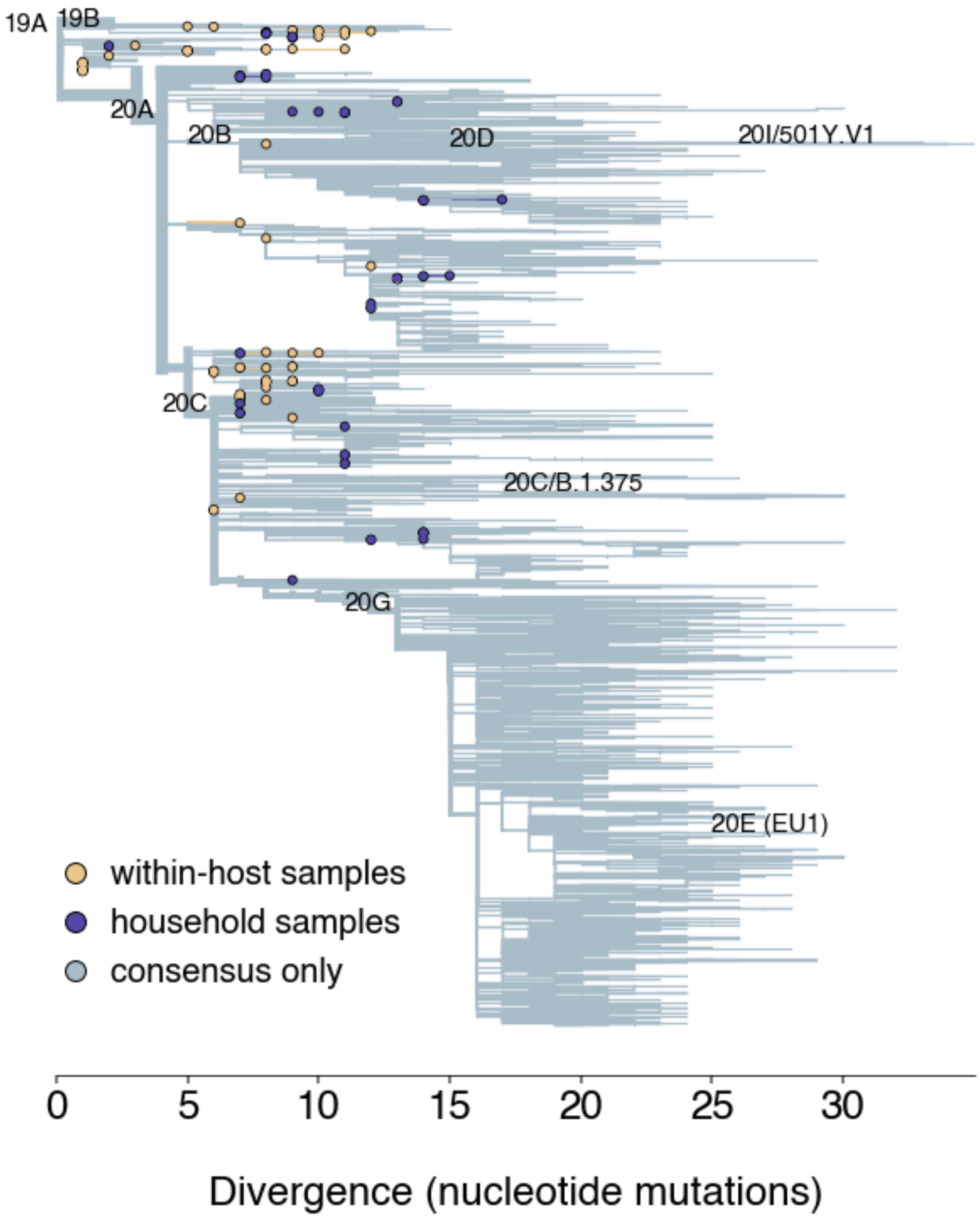


**Supplemental Figure 6: Wisconsin divergence phylogeny**

A full-genome phylogenetic tree built showing X Wisconsin consensus sequences with the Nextstrain pipeline is shown. The x-axis represents divergence expressed as the number of nucleotide mutations. Nextstrain clade labels are shown on the corresponding branch. Yellow tips represent Wisconsin samples that were Illumina sequenced in duplicate and analyzed in this manuscript. Purple tips represent samples from households.

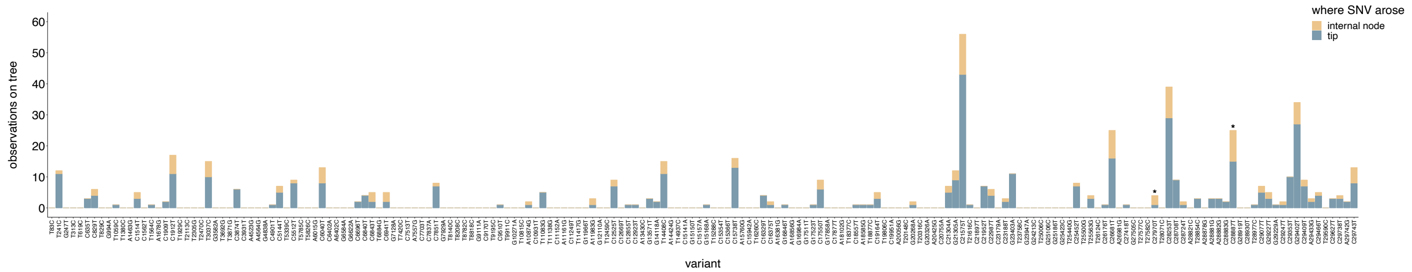


**Supplemental Figure 7: Most iSNVs are not detected on the phylogeny**

We queried every iSNV that was detected within-host (in at least 1 sample) in the global SARS-CoV-2 phylogenetic tree and quantified the number of times that iSNV was detected on an internal node (yellow bar heights) or on a terminal node/tip (blue bar heights). Only approximately 1/3 of all SNVs detected within-host were found on the tree, and none of the indels detected within-host were detected on the phylogeny. Most SNVs that were detected on the tree were rare, and occurred predominantly on terminal nodes. Please note you will likely need to zoom into this figure to clearly read the labels along the x-axis.

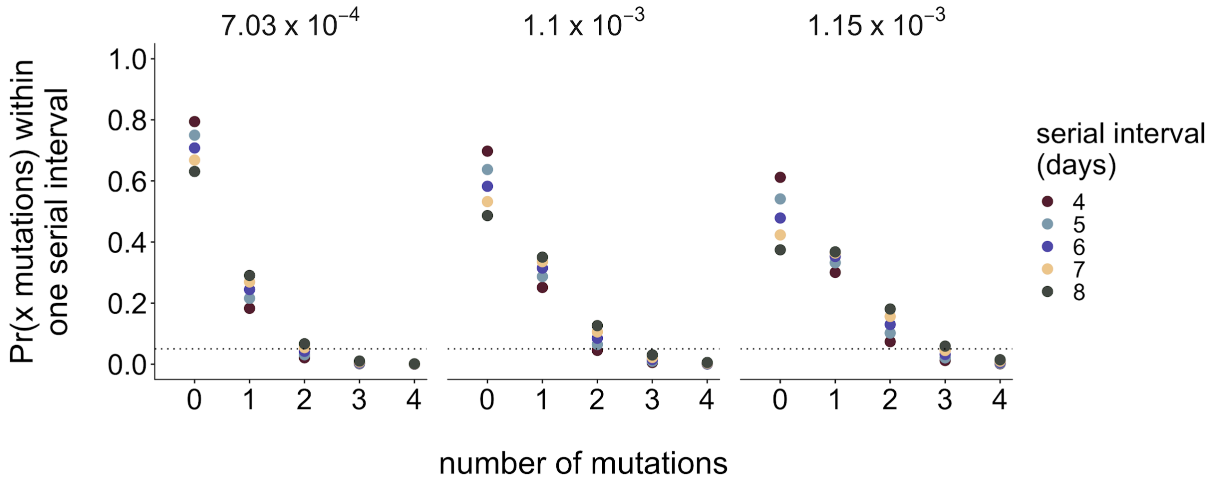


**Supplemental Figure 8: Modeling the expected number of mutations distinguishing genomes separated by one serial interval**

To define whether infections sampled from the same household might be true transmission pairs, we explored the expected number of consensus mutations that should differ between genomes separated by one serial interval. We modeled the probability that 2 consensus genomes will share x mutations as Poisson distributed with lambda equal to the number of mutations expected to accumulate in the SARS-CoV-2 genome over a single serial interval, given a known substitution rate. He et al. estimate a serial interval for SARS-CoV-2 of of 5.8 days, with a 95% confidence interval between 4.8-6.8 days *(35)*. We therefore evaluated serial intervals of 4, 5, 6, 7, and 8 days. For the substitution rate, we use estimates from Duchene et al *(1)*, who estimate a mean substitution rate of $1.10 \times 10^{-3}$ substitutions per site per year, with a 95% credible interval of $7.03 \times 10^{-4}$ and $1.15 \times 10^{-3}$. We evaluated the probabilities that two consensus genomes differ by 0, 1, 2, 3, and 4 mutations given serial intervals ranging from 4-8, and clock rates at the mean, and upper and lower bounds of the 95% credible interval. For each calculated probability, the serial interval is represented by color and the substitution rate is shown above each plot. The dotted line represents a probability of 0.05. Given these combinations of values, the vast majority of consensus genomes are expected to differ by 0-2 mutations.

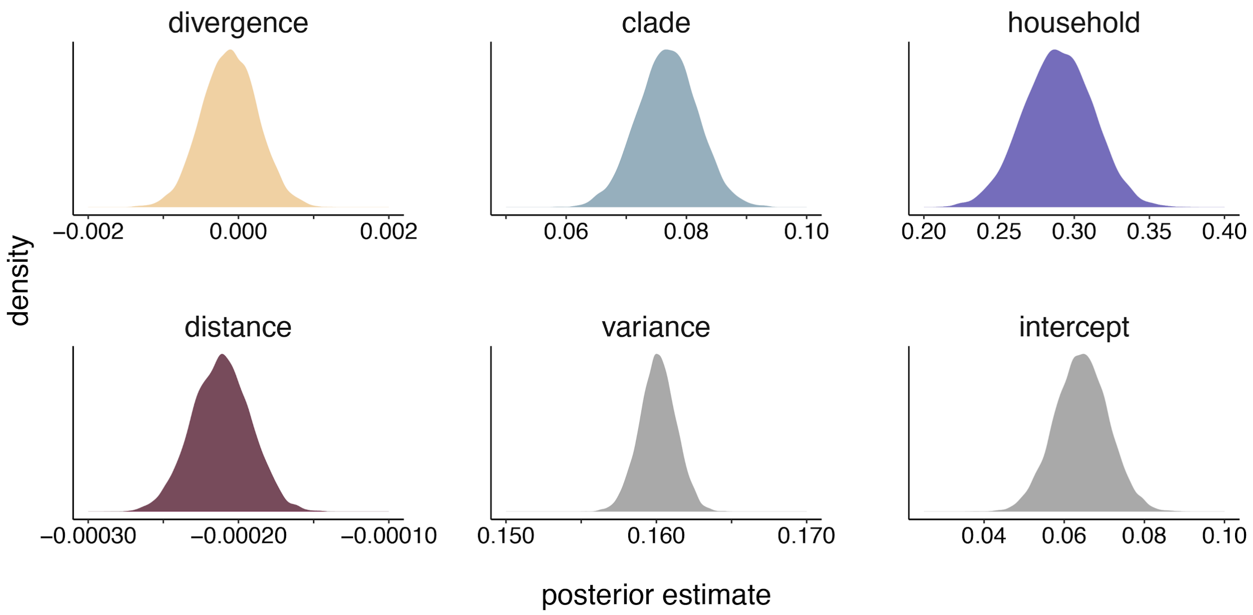


**Supplemental Figure 9: Posterior density estimates for regression coefficients**

For each regression coefficient evaluated in the combined regression model, the full posterior distribution is shown as a density plot. The posterior distribution of the estimated variance and intercept are also shown.

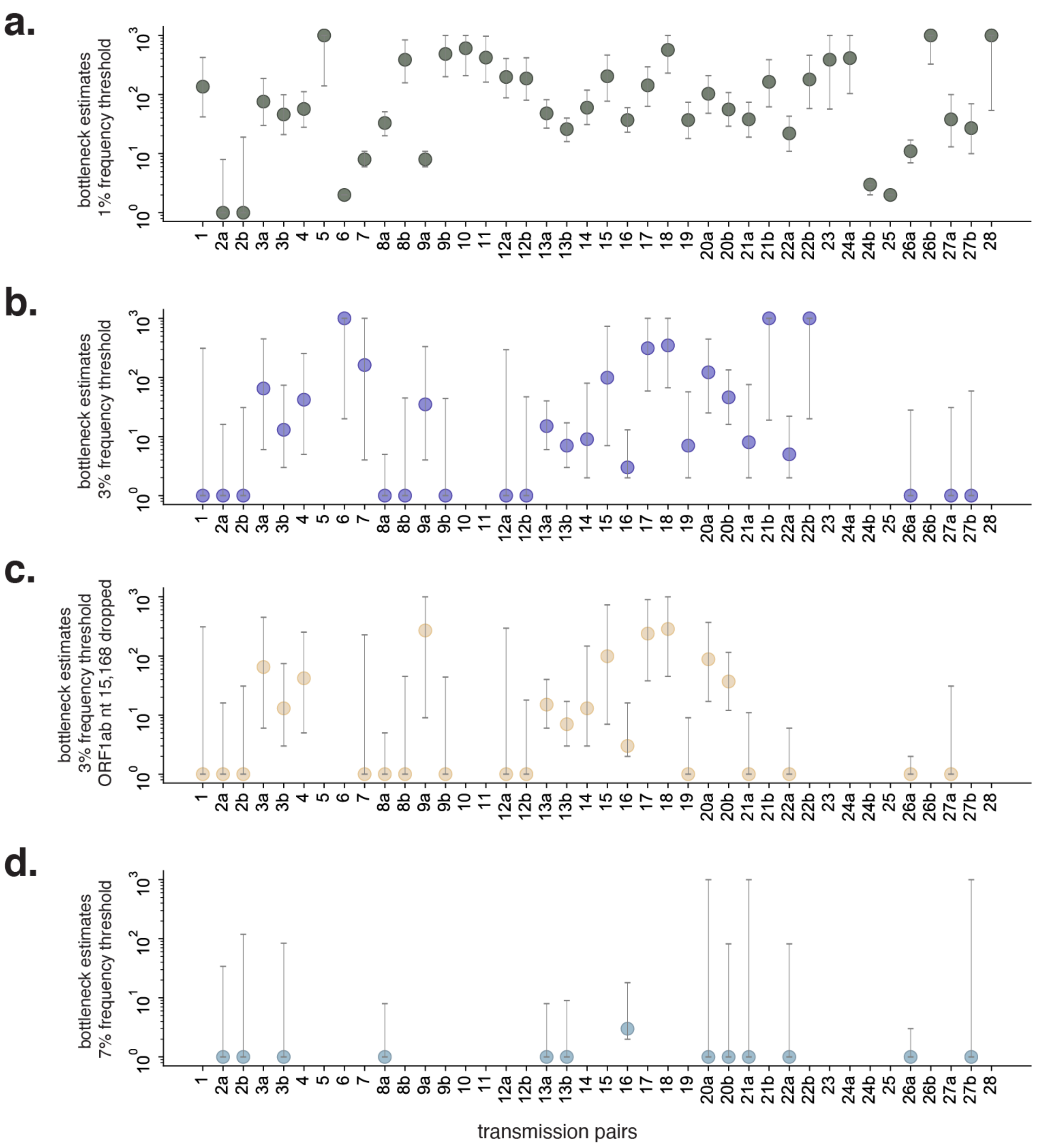


**Supplemental Figure 10: Sensitivity testing of transmission bottleneck estimates**

Maximum likelihood estimates for mean transmission bottleneck size in individual donor-recipient pairs using **a.** 1% frequency threshold, **b.** 3% frequency threshold, **c.** excluding site 15,168 as a possible homoplasy with a 3% frequency threshold, and **d.** 7% frequency threshold. Data are not shown for donor-recipient pairs where no bottleneck estimate could be generated due to lack of variant data. Bidirectional comparisons are indicated with an “a” and “b” following the pair number.

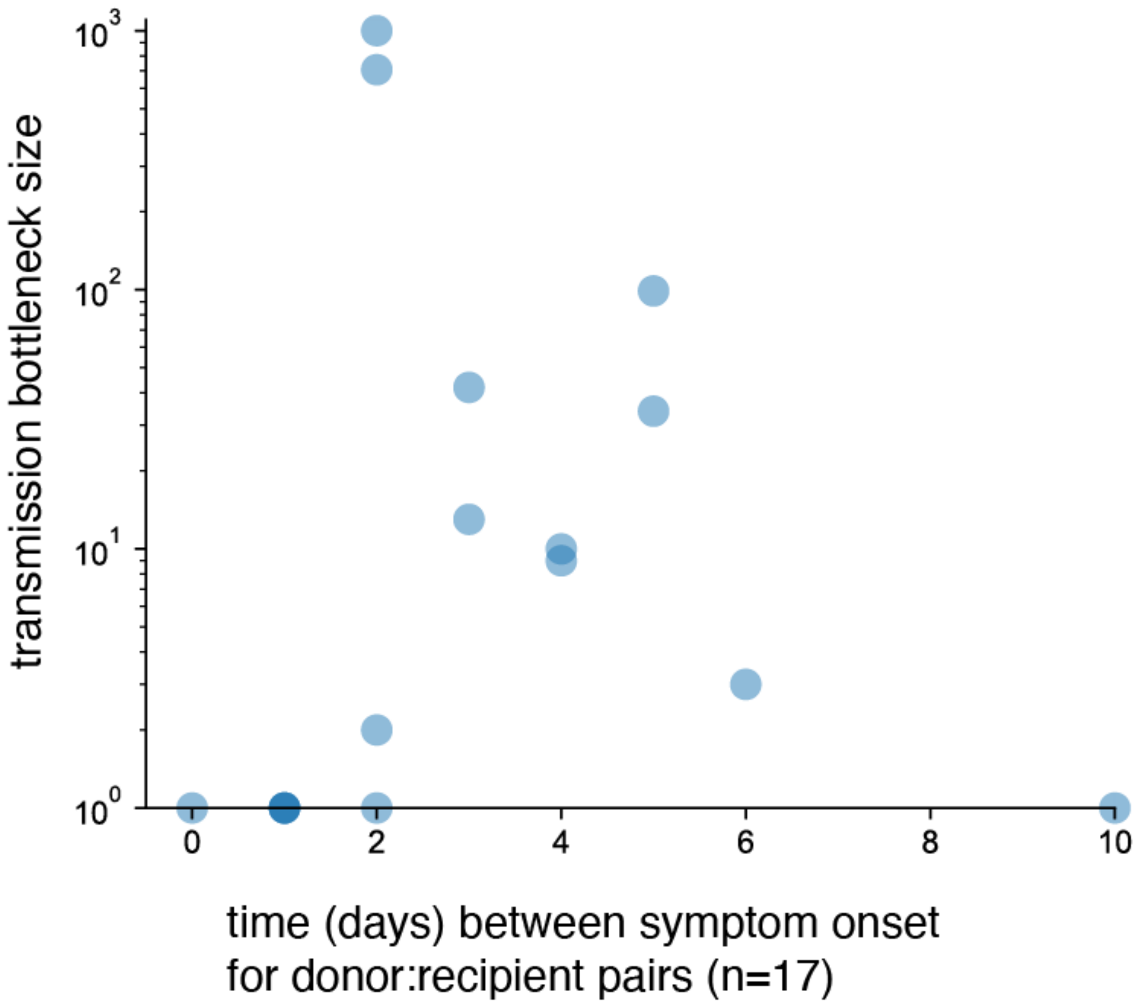


**Supplemental Figure 11: Variance in transmission bottleneck size cannot be explained by time between symptom onset in donor:recipient pairs.** We plotted transmission bottleneck size on the y-axis against time (days) between symptom onset in 17 donor-recipient pairs on the x-axis for which we had symptom metadata.

**Supplemental Table 1**. iSNVs detected in replicate sequencing of the synthetic RNA control (Twist-Biosciences).

| Mutation | | | | | | | | | |
|---|---|---|---|---|---|---|---|---|---|
| **Gene** | **Reference amino acid** | **Amino acid position** | **Variant amino acid** | **Reference nucleotide** | **Nucleotide position** | **Variant nucleotide** | **rep1 percent** | **rep2 percent** | **Average percent** |
| orf1ab | Ser | 1029 | Cys | A | 3350 | T | 0.0406 | 0.0441 | 0.04235 |
| orf1ab | Trp | 2135 | *Stop | G | 6669 | A | 0.0304 | 0.0347 | 0.03255 |
| orf1ab | Gly | 2863 | Val | G | 8853 | T | 0.0103 | 0.011 | 0.01065 |
| orf1ab | Thr | 2967 | Ser | A | 9164 | T | 0.0125 | 0.0109 | 0.0117 |
| M | Leu | 90 | *Stop | T | 26791 | A | 0.1329 | 0.1368 | 0.13485 |
| M | Met | 90 | Val | A | 26793 | G | 0.1313 | 0.1354 | 0.13335 |
| M | Trp | 92 | Arg | T | 26796 | A | 0.131 | 0.1352 | 0.1331 |

**Supplemental Table 2**. Sample identifiers and accession numbers. This table includes strain name, tube/filename, state of collection, county of collection, collection date, GISAID accession number, Genbank accession number, as well as Ct values and RLU values where available for each sample included in this study.

| | | | | | | | Nanopore data | Illumina data | | | |
|---|---|---|---|---|---|---|---|---|---|---|---|
| **Strain** | **Tube** | **State** | **County** | **Collection Date** | **GISAID Accession** | **Genbank Accession** | **BioProject** | **BioProject** | **N1 Ct value** | **N2 Ct value** | **RLU** |
| USA/WI-UW-06/2020 | A | Wisconsin | Dane County | 2020-3-21 | EPI_ISL_417200 | MT772088 | PRJNA614504 | PRJNA718341 | 26.53 | 27.29 | - |
| USA/WI-UW-07/2020 | B | Wisconsin | Dane County | 2020-3-21 | EPI_ISL_417201 | MT772089 | PRJNA614504 | PRJNA718341 | 16.28 | 16.49 | - |
| USA/WI-UW-11/2020 | 1P | Wisconsin | Dane County | 2020-3-15 | EPI_ISL_417505 | MT706133 | PRJNA614504 | PRJNA718341 | - | - | - |
| USA/WI-UW-29/2020 | 5 | Wisconsin | Dane County | 2020-3-24 | EPI_ISL_421287 | MT706150 | PRJNA614504 | PRJNA718341 | 16.14 | 16.05 | - |
| USA/WI-UW-30/2020 | 6 | Wisconsin | Columbia County | 2020-3-26 | EPI_ISL_421288 | MT706151 | PRJNA614504 | PRJNA718341 | 24.76 | 25.41 | - |
| USA/WI-UW-14/2020 | 6P | Wisconsin | Dane County | 2020-3-16 | EPI_ISL_417513 | MT706136 | PRJNA614504 | PRJNA718341 | - | - | - |
| USA/WI-UW-32/2020 | 8 | Wisconsin | Dane County | 2020-3-24 | EPI_ISL_421290 | MT706153 | PRJNA614504 | PRJNA718341 | 24.27 | 24.83 | - |
| USA/WI-UW-34/2020 | 12 | Wisconsin | Dane County | 2020-3-26 | EPI_ISL_421292 | MT706155 | PRJNA614504 | PRJNA718341 | 27.81 | 29.4 | - |
| USA/WI-UW-17/2020 | 12P | Wisconsin | Dane County | 2020-3-13 | EPI_ISL_417517 | MT706139 | PRJNA614504 | PRJNA718341 | - | - | - |
| USA/WI-UW-38/2020 | 17 | Wisconsin | Dane County | 2020-3-25 | EPI_ISL_421296 | MT706159 | PRJNA614504 | PRJNA718341 | 27.98 | 29.23 | - |
| USA/WI-UW-40/2020 | 19 | Wisconsin | Dane County | 2020-3-24 | EPI_ISL_421298 | MT706161 | PRJNA614504 | PRJNA718341 | 23.52 | 24.46 | - |
| USA/WI-UW-39/2020 | 18 | Wisconsin | Dane County | 2020-3-22 | EPI_ISL_421297 | MT706160 | PRJNA614504 | PRJNA718341 | 18.2 | 20.06 | - |
| USA/WI-UW-41/2020 | 20 | Wisconsin | Dane County | 2020-3-25 | EPI_ISL_421299 | MT706162 | PRJNA614504 | PRJNA718341 | 24.32 | 25.31 | - |
| USA/WI-UW-21/2020 | 24P | Wisconsin | Dane County | 2020-3-16 | EPI_ISL_417508 | MT706142 | PRJNA614504 | PRJNA718341 | - | - | - |
| USA/WI-UW-45/2020 | 25 | Wisconsin | Dane County | 2020-3-22 | EPI_ISL_421303 | MT706166 | PRJNA614504 | PRJNA718341 | 20.22 | 20.7 | - |
| USA/WI-UW-22/2020 | 26P | Wisconsin | Dane County | 2020-3-13 | EPI_ISL_417514 | MT706143 | PRJNA614504 | PRJNA718341 | - | - | - |
| USA/WI-UW-48/2020 | 28 | Wisconsin | Dane County | 2020-3-25 | EPI_ISL_421306 | MT706169 | PRJNA614504 | PRJNA718341 | 23.02 | 23.79 | - |
| USA/WI-UW-24/2020 | 29P | Wisconsin | Dane County | 2020-3-15 | EPI_ISL_417512 | MT706145 | PRJNA614504 | PRJNA718341 | - | - | - |
| USA/WI-UW-50/2020 | 30 | Wisconsin | Green County | 2020-3-25 | EPI_ISL_421308 | MT706171 | PRJNA614504 | PRJNA718341 | 19.13 | 19.78 | - |
| USA/WI-UW-51/2020 | 31 | Wisconsin | Dane County | 2020-3-20 | EPI_ISL_421309 | MT706172 | PRJNA614504 | PRJNA718341 | 17.11 | 17.3 | - |

| | | | | | | | | | | | |
|---|---|---|---|---|---|---|---|---|---|---|---|
| USA/WI-UW-52/2020 | 32 | Wisconsin | Dane County | 2020-3-18 | EPI_ISL_421310 | MT706173 | PRJNA614504 | PRJNA718341 | 15.98 | 16.57 | - |
| USA/WI-UW-61/2020 | 44 | Wisconsin | Dane County | 2020-3-23 | EPI_ISL_421319 | MT706182 | PRJNA614504 | PRJNA718341 | 23.28 | 24.11 | - |
| USA/WI-UW-63/2020 | 46 | Wisconsin | Dane County | 2020-3-24 | EPI_ISL_421321 | MT706184 | PRJNA614504 | PRJNA718341 | 24.54 | 25.21 | - |
| USA/WI-UW-65/2020 | 50 | Wisconsin | Dane County | 2020-3-22 | EPI_ISL_421323 | MT706186 | PRJNA614504 | PRJNA718341 | 15.54 | 15.33 | - |
| USA/WI-UW-66/2020 | 51 | Wisconsin | Dane County | 2020-3-24 | EPI_ISL_421324 | MT706187 | PRJNA614504 | PRJNA718341 | 26.52 | 27.61 | - |
| USA/WI-UW-67/2020 | 53 | Wisconsin | Dane County | 2020-3-25 | EPI_ISL_421325 | MT706188 | PRJNA614504 | PRJNA718341 | 25.62 | 27.01 | - |
| USA/WI-UW-68/2020 | 54 | Wisconsin | Dane County | 2020-3-24 | EPI_ISL_421326 | MT706189 | PRJNA614504 | PRJNA718341 | 15.96 | 16.13 | - |
| USA/WI-UW-69/2020 | 55 | Wisconsin | Dane County | 2020-3-19 | EPI_ISL_421327 | MT706190 | PRJNA614504 | PRJNA718341 | 15.83 | 16.07 | - |
| USA/WI-UW-70/2020 | 56 | Wisconsin | Dane County | 2020-3-19 | EPI_ISL_421328 | MT706191 | PRJNA614504 | PRJNA718341 | 20.12 | 20.77 | - |
| USA/WI-UW-71/2020 | 57 | Wisconsin | Dane County | 2020-3-24 | EPI_ISL_421329 | MT706192 | PRJNA614504 | PRJNA718341 | 18.93 | 18.64 | - |
| USA/WI-UW-73/2020 | 60 | Wisconsin | Dane County | 2020-3-24 | EPI_ISL_421331 | MT706194 | PRJNA614504 | PRJNA718341 | 23.69 | 24.93 | - |
| USA/WI-UW-74/2020 | 61 | Wisconsin | Dane County | 2020-3-20 | EPI_ISL_421332 | MT706195 | PRJNA614504 | PRJNA718341 | 14.19 | 14.36 | - |
| USA/WI-UW-76/2020 | 64 | Wisconsin | Dane County | 2020-3-22 | EPI_ISL_421334 | MT706197 | PRJNA614504 | PRJNA718341 | 17.49 | 17.59 | - |
| USA/WI-UW-77/2020 | 65 | Wisconsin | Dane County | 2020-3-19 | EPI_ISL_421335 | MT706198 | PRJNA614504 | PRJNA718341 | 20.19 | 20.65 | - |
| USA/WI-UW-84/2020 | 74 | Wisconsin | Dane County | 2020-3-24 | EPI_ISL_421343 | MT706205 | PRJNA614504 | PRJNA718341 | 23.12 | 23.82 | - |
| USA/WI-UW-85/2020 | 79 | Wisconsin | Dane County | 2020-4-2 | EPI_ISL_425142 | MT706206 | PRJNA614504 | PRJNA718341 | 24.4 | - | - |
| USA/WI-UW-86/2020 | 80 | Wisconsin | Dane County | 2020-4-2 | EPI_ISL_425143 | MT706207 | PRJNA614504 | PRJNA718341 | 22.1 | - | - |
| USA/WI-UW-87/2020 | 81 | Wisconsin | Dane County | 2020-4-2 | EPI_ISL_425144 | MT706208 | PRJNA614504 | PRJNA718341 | 22.1 | - | - |
| USA/WI-UW-88/2020 | 82 | Wisconsin | Dane County | 2020-4-5 | EPI_ISL_425145 | MT706209 | PRJNA614504 | PRJNA718341 | 25.29 | 25.93 | - |
| USA/WI-UW-96/2020 | 94 | Wisconsin | Dane County | 2020-4-1 | EPI_ISL_425153 | MT706216 | PRJNA614504 | PRJNA718341 | 17.33 | 18.05 | - |
| USA/WI-UW-97/2020 | 95 | Wisconsin | Dane County | 2020-3-30 | EPI_ISL_425154 | MT706217 | PRJNA614504 | PRJNA718341 | 27.3 | - | - |
| USA/WI-UW-99/2020 | 99 | Wisconsin | Dane County | 2020-4-2 | EPI_ISL_425156 | MT706219 | PRJNA614504 | PRJNA718341 | 18.8 | - | - |

| | | | | | | | | | | | |
|---|---|---|---|---|---|---|---|---|---|---|---|
| USA/WI-UW-110/2020 | 117 | Wisconsin | Dane County | 2020-3-31 | EPI_ISL_425167 | MT706230 | PRJNA614504 | PRJNA718341 | 22.2 | - | - |
| USA/WI-UW-111/2020 | 118 | Wisconsin | Dane County | 2020-3-31 | EPI_ISL_425168 | MT706231 | PRJNA614504 | PRJNA718341 | 25.61 | 26.29 | - |
| USA/WI-UW-116/2020 | 124 | Wisconsin | Dane County | 2020-3-30 | EPI_ISL_425173 | MT706236 | PRJNA614504 | PRJNA718341 | 31.81 | 33.31 | - |
| USA/WI-UW-117/2020 | 125 | Wisconsin | Dane County | 2020-3-30 | EPI_ISL_425174 | MT706237 | PRJNA614504 | PRJNA718341 | 28.2 | - | - |
| USA/WI-UW-119/2020 | 128 | Wisconsin | Dane County | 2020-4-10 | EPI_ISL_425176 | MT706239 | PRJNA614504 | PRJNA718341 | 14.76 | 14.82 | - |
| USA/WI-UW-120/2020 | 130 | Wisconsin | Dane County | 2020-4-13 | EPI_ISL_427427 | MT706240 | PRJNA614504 | PRJNA718341 | 18.3 | - | - |
| USA/WI-UW-255/2020 | 139 | Wisconsin | Dane County | 2020-4-2 | EPI_ISL_428729 | MT706248 | PRJNA614504 | PRJNA718341 | - | - | - |
| USA/WI-UW-124/2020 | 145 | Wisconsin | Dane County | 2020-4-7 | EPI_ISL_427431 | MT706252 | PRJNA614504 | PRJNA718341 | 17.5 | - | - |
| USA/WI-UW-127/2020 | 148 | Wisconsin | Dane County | 2020-4-7 | EPI_ISL_427434 | MT706255 | PRJNA614504 | PRJNA718341 | 19.1 | - | - |
| USA/WI-UW-129/2020 | 151 | Wisconsin | Dane County | 2020-4-6 | EPI_ISL_427436 | MT706257 | PRJNA614504 | PRJNA718341 | 30.7 | - | - |
| USA/WI-UW-132/2020 | 158 | Wisconsin | Rock County | 2020-4-10 | EPI_ISL_427439 | MT706260 | PRJNA614504 | PRJNA718341 | 17.1 | - | - |
| USA/WI-UW-140/2020 | 168 | Wisconsin | Dane County | 2020-4-9 | EPI_ISL_427447 | MT706268 | PRJNA614504 | PRJNA718341 | 21.43 | - | - |
| USA/WI-UW-144/2020 | 172 | Wisconsin | Dane County | 2020-4-6 | EPI_ISL_427451 | MT706272 | PRJNA614504 | PRJNA718341 | 23.92 | 24.31 | - |
| USA/WI-UW-146/2020 | 175 | Wisconsin | Dane County | 2020-4-6 | EPI_ISL_427453 | MT706274 | PRJNA614504 | PRJNA718341 | 26.1 | - | - |
| USA/IL-UW-149/2020 | 182 | Illinois | Winnebago County | 2020-4-7 | EPI_ISL_427456 | MT706277 | PRJNA614504 | PRJNA718341 | 12.6 | - | - |
| USA/WI-UW-154/2020 | 188 | Wisconsin | Monroe County | 2020-4-12 | EPI_ISL_427461 | MT706282 | PRJNA614504 | PRJNA718341 | 21.5 | - | - |
| USA/WI-UW-158/2020 | 195 | Wisconsin | Milwaukee County | 2020-3-15 | EPI_ISL_428254 | MT706286 | PRJNA614504 | PRJNA718341 | 23.93 | 24.33 | - |
| USA/WI-UW-159/2020 | 196 | Wisconsin | Milwaukee County | 2020-3-15 | EPI_ISL_428255 | MT706287 | PRJNA614504 | PRJNA718341 | 20.57 | 21.06 | - |
| USA/WI-UW-160/2020 | 197 | Wisconsin | Milwaukee County | 2020-3-15 | EPI_ISL_428256 | MT706288 | PRJNA614504 | PRJNA718341 | 18.6 | 19.08 | - |
| USA/WI-UW-179/2020 | 218 | Wisconsin | Milwaukee County | 2020-3-21 | EPI_ISL_428275 | MT706307 | PRJNA614504 | PRJNA718341 | 19.89 | 18.58 | - |
| USA/WI-UW-188/2020 | 229 | Wisconsin | Milwaukee County | 2020-3-23 | EPI_ISL_428284 | MT706316 | PRJNA614504 | PRJNA718341 | 30.05 | 29.06 | - |
| USA/WI-UW-201/2020 | 245 | Wisconsin | Ozaukee County | 2020-3-25 | EPI_ISL_428297 | MT706329 | PRJNA614504 | PRJNA718341 | 21.19 | 21.23 | - |

| | | | | | | | | | | | |
|---|---|---|---|---|---|---|---|---|---|---|---|
| USA/WI-UW-205/2020 | 249 | Wisconsin | Milwaukee County | 2020-3-25 | EPI_ISL_428301 | MT706333 | PRJNA614504 | PRJNA718341 | 27.03 | 27.23 | - |
| USA/WI-UW-208/2020 | 252 | Wisconsin | Milwaukee County | 2020-3-25 | EPI_ISL_428304 | MT706336 | PRJNA614504 | PRJNA718341 | 25.72 | 24.87 | - |
| USA/WI-UW-211/2020 | 255 | Wisconsin | Milwaukee County | 2020-3-25 | EPI_ISL_428307 | MT706339 | PRJNA614504 | PRJNA718341 | 26.72 | 27.21 | - |
| USA/WI-UW-214/2020 | 258 | Wisconsin | Ozaukee County | 2020-3-25 | EPI_ISL_428310 | MT706342 | PRJNA614504 | PRJNA718341 | 21.56 | 21.52 | - |
| USA/WI-UW-217/2020 | 261 | Wisconsin | Milwaukee County | 2020-3-25 | EPI_ISL_428313 | MT706345 | PRJNA614504 | PRJNA718341 | 29.14 | 29.6 | - |
| USA/WI-UW-223/2020 | 268 | Wisconsin | Milwaukee County | 2020-3-26 | EPI_ISL_428319 | MT706351 | PRJNA614504 | PRJNA718341 | 26.67 | 28.25 | - |
| USA/WI-UW-225/2020 | 270 | Wisconsin | Milwaukee County | 2020-3-26 | EPI_ISL_428321 | MT706353 | PRJNA614504 | PRJNA718341 | 18.55 | 18.91 | - |
| USA/WI-UW-230/2020 | 275 | Wisconsin | Milwaukee County | 2020-3-26 | EPI_ISL_428326 | MT706358 | PRJNA614504 | PRJNA718341 | 24.4 | 25.99 | - |
| USA/WI-UW-231/2020 | 276 | Wisconsin | Milwaukee County | 2020-3-26 | EPI_ISL_428327 | MT706359 | PRJNA614504 | PRJNA718341 | 19.42 | 19.94 | - |
| USA/WI-UW-232/2020 | 277 | Wisconsin | Milwaukee County | 2020-3-27 | EPI_ISL_428328 | MT706360 | PRJNA614504 | PRJNA718341 | 19.54 | 18.33 | - |
| USA/WI-UW-238/2020 | 283 | Wisconsin | Milwaukee County | 2020-3-27 | EPI_ISL_428334 | MT706366 | PRJNA614504 | PRJNA718341 | 26.38 | 25.52 | - |
| USA/WI-UW-242/2020 | 288 | Wisconsin | Milwaukee County | 2020-3-28 | EPI_ISL_428338 | MT706370 | PRJNA614504 | PRJNA718341 | 17.93 | 17.52 | - |
| USA/WI-UW-243/2020 | 289 | Wisconsin | Ozaukee County | 2020-3-28 | EPI_ISL_428339 | MT706371 | PRJNA614504 | PRJNA718341 | 25.29 | 25.04 | - |
| USA/WI-UW-244/2020 | 290 | Wisconsin | Ozaukee County | 2020-3-28 | EPI_ISL_428340 | MT706372 | PRJNA614504 | PRJNA718341 | 19.65 | 19.25 | - |
| USA/WI-UW-246/2020 | 292 | Wisconsin | Milwaukee County | 2020-3-28 | EPI_ISL_428342 | - | PRJNA614504 | PRJNA718341 | 22.16 | 21.97 | - |
| Tube-302 | 302 | Wisconsin | Dane County | 2020-4-16 | - | - | - | PRJNA718341 | 29.5 | - | - |
| Tube-303 | 303 | Wisconsin | Monroe County | 2020-4-16 | - | - | - | PRJNA718341 | - | - | - |
| Tube-304 | 304 | Wisconsin | Dane County | 2020-4-19 | - | - | - | PRJNA718341 | 30.7 | - | - |
| USA/WI-UW-348/2020 | 310 | Wisconsin | Dane County | 2020-4-15 | EPI_ISL_450702 | MT506887 | PRJNA614504 | PRJNA718341 | - | - | - |
| USA/WI-UW-351/2020 | 316 | Wisconsin | Dane County | 2020-4-14 | EPI_ISL_450705 | MT506890 | PRJNA614504 | PRJNA718341 | 25.51 | 26.64 | - |
| USA/WI-UW-367/2020 | 338 | Wisconsin | Dane County | 2020-4-1 | EPI_ISL_450721 | MT506906 | PRJNA614504 | PRJNA718341 | - | - | - |
| USA/WI-UW-273/2020 | 356 | Wisconsin | Milwaukee County | 2020-3-31 | EPI_ISL_436567 | MT706381 | PRJNA614504 | PRJNA718341 | 20.29 | 20.47 | - |

| | | | | | | | | | | | |
|---|---|---|---|---|---|---|---|---|---|---|---|
| USA/WI-UW-275/2020 | 358 | Wisconsin | Milwaukee County | 2020-4-1 | EPI_ISL_436569 | MT706383 | PRJNA614504 | PRJNA718341 | 18.54 | 18.14 | - |
| USA/WI-UW-276/2020 | 359 | Wisconsin | Milwaukee County | 2020-4-1 | EPI_ISL_436570 | MT706384 | PRJNA614504 | PRJNA718341 | 19.27 | 19.02 | - |
| USA/WI-UW-277/2020 | 361 | Wisconsin | Milwaukee County | 2020-4-2 | EPI_ISL_436571 | MT706385 | PRJNA614504 | PRJNA718341 | 20.28 | 20.06 | - |
| USA/WI-UW-278/2020 | 365 | Wisconsin | Milwaukee County | 2020-4-3 | EPI_ISL_436572 | MT706386 | PRJNA614504 | PRJNA718341 | 16.08 | 16.12 | - |
| USA/WI-UW-279/2020 | 366 | Wisconsin | Milwaukee County | 2020-4-3 | EPI_ISL_436573 | MT706387 | PRJNA614504 | PRJNA718341 | 15.6 | 15.35 | - |
| USA/WI-UW-282/2020 | 370 | Wisconsin | Milwaukee County | 2020-4-6 | EPI_ISL_436576 | MT706390 | PRJNA614504 | PRJNA718341 | 15.39 | 14.91 | - |
| USA/WI-UW-285/2020 | 374 | Wisconsin | Milwaukee County | 2020-4-6 | EPI_ISL_436579 | MT706393 | PRJNA614504 | PRJNA718341 | 27.55 | 27.17 | - |
| USA/WI-UW-286/2020 | 376 | Wisconsin | Milwaukee County | 2020-4-6 | EPI_ISL_436580 | MT706394 | PRJNA614504 | PRJNA718341 | 25.2 | 25.09 | - |
| USA/WI-UW-287/2020 | 377 | Wisconsin | Milwaukee County | 2020-4-6 | EPI_ISL_436581 | MT706395 | PRJNA614504 | PRJNA718341 | 23.43 | 23.35 | - |
| USA/WI-UW-296/2020 | 388 | Wisconsin | Milwaukee County | 2020-4-9 | EPI_ISL_436590 | MT706404 | PRJNA614504 | PRJNA718341 | 16.13 | 15.52 | - |
| USA/WI-UW-299/2020 | 391 | Wisconsin | Milwaukee County | 2020-4-13 | EPI_ISL_436593 | MT706407 | PRJNA614504 | PRJNA718341 | 18.84 | 18.37 | - |
| USA/WI-UW-302/2020 | 397 | Wisconsin | Milwaukee County | 2020-4-13 | EPI_ISL_436596 | MT706410 | PRJNA614504 | PRJNA718341 | 17.6 | 17.08 | - |
| USA/WI-UW-306/2020 | 402 | Wisconsin | Milwaukee County | 2020-4-14 | EPI_ISL_436600 | MT706414 | PRJNA614504 | PRJNA718341 | 30.42 | 31.22 | - |
| USA/WI-UW-310/2020 | 408 | Wisconsin | Milwaukee County | 2020-4-15 | EPI_ISL_436604 | MT706418 | PRJNA614504 | PRJNA718341 | 20.64 | 19.45 | - |
| USA/WI-UW-315/2020 | 418 | Wisconsin | Milwaukee County | 2020-4-17 | EPI_ISL_436609 | MT706423 | PRJNA614504 | PRJNA718341 | 26.33 | 26.42 | - |
| USA/WI-UW-323/2020 | 437 | Wisconsin | Milwaukee County | 2020-4-23 | EPI_ISL_436617 | MT706430 | PRJNA614504 | PRJNA718341 | 26.2 | 28.6 | - |
| USA/WI-UW-333/2020 | 453 | Wisconsin | Milwaukee County | 2020-3-24 | EPI_ISL_436627 | MT706439 | PRJNA614504 | PRJNA718341 | 27.45 | 26.54 | - |
| USA/WI-UW-334/2020 | 454 | Wisconsin | Milwaukee County | 2020-3-24 | EPI_ISL_436628 | MT706440 | PRJNA614504 | PRJNA718341 | 23.24 | 22.79 | - |
| USA/WI-UW-337/2020 | 461 | Wisconsin | Milwaukee County | 2020-3-26 | EPI_ISL_436631 | MT706443 | PRJNA614504 | PRJNA718341 | 26.46 | 26.5 | - |
| USA/WI-UW-338/2020 | 462 | Wisconsin | Milwaukee County | 2020-3-26 | EPI_ISL_436632 | MT706444 | PRJNA614504 | PRJNA718341 | 27.72 | 27.6 | - |
| USA/WI-UW-340/2020 | 468 | Wisconsin | Milwaukee County | 2020-4-1 | EPI_ISL_436634 | MT706446 | PRJNA614504 | PRJNA718341 | 35.15 | 33.47 | - |
| USA/WI-UW-341/2020 | 469 | Wisconsin | Milwaukee County | 2020-4-2 | EPI_ISL_436635 | MT706447 | PRJNA614504 | PRJNA718341 | 23.18 | 22.53 | - |

| | | | | | | | | | | | |
|---|---|---|---|---|---|---|---|---|---|---|---|
| USA/WI-UW-389/2020 | 552 | Wisconsin | Dane County | 2020-5-26 | EPI_ISL_480371 | MT772540 | PRJNA614504 | PRJNA718341 | 23.2 | - | - |
| USA/WI-UW-391/2020 | 554 | Wisconsin | Dane County | 2020-5-26 | EPI_ISL_480373 | MT772542 | PRJNA614504 | PRJNA718341 | 17.7 | - | - |
| USA/WI-UW-432/2020 | 738 | Wisconsin | Dane County | 2020-6-17 | EPI_ISL_484807 | MT750020 | PRJNA614504 | PRJNA718341 | 21.6 | - | - |
| USA/WI-UW-438/2020 | 744 | Wisconsin | Dane County | 2020-6-15 | EPI_ISL_484813 | MT750026 | PRJNA614504 | PRJNA718341 | 19.4 | - | - |
| USA/WI-UW-443/2020 | 749 | Wisconsin | Dane County | 2020-6-19 | EPI_ISL_484818 | MT750030 | PRJNA614504 | PRJNA718341 | 19.4 | - | - |
| USA/WI-UW-476/2020 | 783 | Wisconsin | Dane County | 2020-6-12 | EPI_ISL_484851 | MT750060 | PRJNA614504 | PRJNA718341 | 20.6 | - | - |
| USA/WI-UW-536/2020 | 849 | Wisconsin | Dane County | 2020-6-24 | EPI_ISL_484911 | MT750116 | PRJNA614504 | PRJNA718341 | - | - | 1211 |
| USA/WI-UW-544/2020 | 884 | Wisconsin | Dane County | 2020-6-22 | EPI_ISL_484919 | MT750124 | PRJNA614504 | PRJNA718341 | - | - | 1334 |
| USA/WI-UW-546/2020 | 887 | Wisconsin | Dane County | 2020-6-19 | EPI_ISL_484921 | MT750126 | PRJNA614504 | PRJNA718341 | - | - | 1282 |
| USA/WI-UW-551/2020 | 893 | Wisconsin | Dane County | 2020-6-23 | EPI_ISL_484926 | MT750131 | PRJNA614504 | PRJNA718341 | - | - | 1254 |
| USA/WI-UW-575/2020 | 903 | Wisconsin | Dane County | 2020-6-23 | EPI_ISL_484950 | MT750154 | PRJNA614504 | PRJNA718341 | - | - | 1238 |
| USA/WI-UW-577/2020 | 906 | Wisconsin | Dane County | 2020-6-24 | EPI_ISL_484952 | MT750156 | PRJNA614504 | PRJNA718341 | - | - | 1288 |
| USA/WI-UW-586/2020 | 916 | Wisconsin | Dane County | 2020-6-19 | EPI_ISL_484961 | MT750165 | PRJNA614504 | PRJNA718341 | - | - | 1274 |
| USA/WI-UW-598/2020 | 956 | Wisconsin | Dane County | 2020-6-27 | EPI_ISL_484973 | MT750176 | PRJNA614504 | PRJNA718341 | - | - | 1240 |
| USA/WI-UW-601/2020 | 961 | Wisconsin | Dane County | 2020-6-25 | EPI_ISL_484976 | MT750179 | PRJNA614504 | PRJNA718341 | - | - | 1155 |
| USA/WI-UW-602/2020 | 962 | Wisconsin | Dane County | 2020-6-28 | EPI_ISL_484977 | - | PRJNA614504 | PRJNA718341 | - | - | 1203 |
| USA/WI-UW-689/2020 | 1049 | Wisconsin | Dane County | 2020-6-28 | EPI_ISL_491369 | MT772466 | PRJNA614504 | PRJNA718341 | - | - | 1217 |
| USA/WI-UW-694/2020 | 1061 | Wisconsin | Dane County | 2020-6-30 | EPI_ISL_491372 | - | PRJNA614504 | PRJNA718341 | - | - | 1218 |
| USA/WI-UW-697/2020 | 1064 | Wisconsin | Dane County | 2020-7-2 | EPI_ISL_491375 | MT772473 | PRJNA614504 | PRJNA718341 | - | - | 1257 |
| USA/WI-UW-721/2020 | 1103 | Wisconsin | Dane County | 2020-7-1 | EPI_ISL_491396 | - | PRJNA614504 | PRJNA718341 | - | - | 1177 |
| USA/WI-UW-722/2020 | 1104 | Wisconsin | Dane County | 2020-6-30 | EPI_ISL_491397 | - | PRJNA614504 | PRJNA718341 | - | - | 1276 |
| USA/WI-UW-747/2020 | 1144 | Wisconsin | Dane County | 2020-7-2 | EPI_ISL_491420 | MT772518 | PRJNA614504 | PRJNA718341 | - | - | 1222 |

| | | | | | | | | | | | |
|---|---|---|---|---|---|---|---|---|---|---|---|
| USA/WI-UW-749/2020 | 1147 | Wisconsin | Dane County | 2020-6-30 | EPI_ISL_491422 | - | PRJNA614504 | PRJNA718341 | - | - | 1287 |
| USA/WI-UW-756/2020 | 1157 | Wisconsin | Dane County | 2020-7-5 | EPI_ISL_495461 | MT795871 | PRJNA614504 | PRJNA718341 | - | - | 1233 |
| USA/WI-UW-780/2020 | 1195 | Wisconsin | Dane County | 2020-7-3 | EPI_ISL_495484 | MT795891 | PRJNA614504 | PRJNA718341 | - | - | 1269 |
| USA/WI-UW-784/2020 | 1199 | Wisconsin | Dane County | 2020-7-6 | EPI_ISL_495488 | - | PRJNA614504 | PRJNA718341 | - | - | 1238 |
| USA/WI-UW-798/2020 | 1217 | Wisconsin | Dane County | 2020-7-6 | EPI_ISL_495502 | - | PRJNA614504 | PRJNA718341 | - | - | 1223 |
| USA/WI-UW-855/2020 | 1282 | Wisconsin | Dane County | 2020-7-9 | EPI_ISL_509861 | MT846545 | PRJNA614504 | PRJNA718341 | - | - | 1247 |
| USA/WI-UW-861/2020 | 1293 | Wisconsin | Dane County | 2020-7-14 | EPI_ISL_509864 | MT846550 | PRJNA614504 | PRJNA718341 | - | - | 1210 |
| USA/WI-UW-863/2020 | 1297 | Wisconsin | Dane County | 2020-7-13 | EPI_ISL_509866 | MT846552 | PRJNA614504 | PRJNA718341 | - | - | 1159 |
| USA/WI-UW-874/2020 | 1326 | Wisconsin | Dane County | 2020-7-13 | EPI_ISL_509876 | MT846562 | PRJNA614504 | PRJNA718341 | - | - | 1227 |
| USA/WI-UW-876/2020 | 1328 | Wisconsin | Dane County | 2020-7-13 | EPI_ISL_509878 | MT846564 | PRJNA614504 | PRJNA718341 | - | - | 1241 |
| USA/WI-UW-893/2020 | 1346 | Wisconsin | Dane County | 2020-7-12 | EPI_ISL_509895 | MT846581 | PRJNA614504 | PRJNA718341 | - | - | 1181 |
| USA/WI-UW-895/2020 | 1353 | Wisconsin | Dane County | 2020-7-13 | EPI_ISL_509897 | MT846583 | PRJNA614504 | PRJNA718341 | - | - | 1200 |
| USA/WI-UW-897/2020 | 1357 | Wisconsin | Dane County | 2020-7-12 | EPI_ISL_509899 | MT846585 | PRJNA614504 | PRJNA718341 | - | - | 1224 |
| USA/WI-UW-906/2020 | 1373 | Wisconsin | Dane County | 2020-7-15 | EPI_ISL_509907 | MT846593 | PRJNA614504 | PRJNA718341 | - | - | 1246 |
| USA/WI-UW-916/2020 | 1388 | Wisconsin | Dane County | 2020-7-12 | EPI_ISL_509917 | MT846603 | PRJNA614504 | PRJNA718341 | - | - | 1209 |
| USA/WI-UW-927/2020 | 1409 | Wisconsin | Dane County | 2020-7-13 | EPI_ISL_509927 | MT846614 | PRJNA614504 | PRJNA718341 | - | - | 1160 |
| USA/WI-UW-931/2020 | 1414 | Wisconsin | Dane County | 2020-7-13 | EPI_ISL_509931 | MT846618 | PRJNA614504 | PRJNA718341 | - | - | 1221 |
| USA/WI-UW-986/2020 | 1495 | Wisconsin | Dane County | 2020-7-16 | EPI_ISL_509982 | MT846672 | PRJNA614504 | PRJNA718341 | - | - | 1267 |
| USA/WI-UW-991/2020 | 1502 | Wisconsin | Dane County | 2020-7-16 | EPI_ISL_509986 | MT846677 | PRJNA614504 | PRJNA718341 | - | - | 1265 |
| USA/WI-UW-997/2020 | 1512 | Wisconsin | Dane County | 2020-7-16 | EPI_ISL_509991 | MT846683 | PRJNA614504 | PRJNA718341 | - | - | 1220 |

**Supplemental Table 3**. ARTIC v3 primer sequences used to amplify cDNA for library preparation.

| name | pool | sequence | length | %gc | tm (use 65) |
|---|---|---|---|---|---|
| nCoV-2019_1_LEFT | nCoV-2019_1 | ACCAACCAACTTTCGATCTCTTGT | 24 | 41.67 | 60.69 |
| nCoV-2019_1_RIGHT | nCoV-2019_1 | CATCTTTAAGATGTTGACGTGCCTC | 25 | 44 | 60.45 |
| nCoV-2019_2_LEFT | nCoV-2019_2 | CTGTTTTACAGGTTCGCGACGT | 22 | 50 | 61.67 |
| nCoV-2019_2_RIGHT | nCoV-2019_2 | TAAGGATCAGTGCCAAGCTCGT | 22 | 50 | 61.74 |
| nCoV-2019_3_LEFT | nCoV-2019_1 | CGGTAATAAAGGAGCTGGTGGC | 22 | 54.55 | 61.32 |
| nCoV-2019_3_RIGHT | nCoV-2019_1 | AAGGTGTCTGCAATTCATAGCTCT | 24 | 41.67 | 60.32 |
| nCoV-2019_4_LEFT | nCoV-2019_2 | GGTGTATACTGCTGCCGTGAAC | 22 | 54.55 | 61.56 |
| nCoV-2019_4_RIGHT | nCoV-2019_2 | CACAAGTAGTGGCACCTTCTTTAGT | 25 | 44 | 60.97 |
| nCoV-2019_5_LEFT | nCoV-2019_1 | TGGTGAAACTTCATGGCAGACG | 22 | 50 | 61.39 |
| nCoV-2019_5_RIGHT | nCoV-2019_1 | ATTGATGTTGACTTTCTCTTTTTGGAGT | 28 | 32.14 | 60.17 |
| nCoV-2019_6_LEFT | nCoV-2019_2 | GGTGTTGTTGGAGAAGGTTCCG | 22 | 54.55 | 61.64 |
| nCoV-2019_6_RIGHT | nCoV-2019_2 | TAGCGGCCTTCTGTAAAACACG | 22 | 50 | 61.18 |
| nCoV-2019_7_LEFT | nCoV-2019_1 | ATCAGAGGCTGCTCGTGTTGTA | 22 | 50 | 61.73 |
| nCoV-2019_7_LEFT_alt0 | nCoV-2019_1 | CATTTGCATCAGAGGCTGCTCG | 22 | 54.55 | 62.44 |
| nCoV-2019_7_RIGHT | nCoV-2019_1 | TGCACAGGTGACAATTTGTCCA | 22 | 45.45 | 60.95 |
| nCoV-2019_7_RIGHT_alt5 | nCoV-2019_1 | AGGTGACAATTTGTCCACCGAC | 22 | 50 | 61.07 |
| nCoV-2019_8_LEFT | nCoV-2019_2 | AGAGTTTCTTAGAGACGGTTGGGA | 24 | 45.83 | 61 |
| nCoV-2019_8_RIGHT | nCoV-2019_2 | GCTTCAACAGCTTCACTAGTAGGT | 24 | 45.83 | 60.56 |
| nCoV-2019_9_LEFT | nCoV-2019_1 | TCCCACAGAAGTGTTAACAGAGGA | 24 | 45.83 | 61.18 |
| nCoV-2019_9_LEFT_alt4 | nCoV-2019_1 | TTCCCACAGAAGTGTTAACAGAGG | 24 | 45.83 | 60.44 |
| nCoV-2019_9_RIGHT | nCoV-2019_1 | ATGACAGCATCTGCCACAACAC | 22 | 50 | 61.71 |
| nCoV-2019_9_RIGHT_alt2 | nCoV-2019_1 | GACAGCATCTGCCACAACACAG | 22 | 54.55 | 62.26 |
| nCoV-2019_10_LEFT | nCoV-2019_2 | TGAGAAGTGCTCTGCCTATACAGT | 24 | 45.83 | 61.12 |
| nCoV-2019_10_RIGHT | nCoV-2019_2 | TCATCTAACCAATCTTCTTCTTGCTCT | 27 | 37.04 | 60.31 |
| nCoV-2019_11_LEFT | nCoV-2019_1 | GGAATTTGGTGCCACTTCTGCT | 22 | 50 | 61.66 |
| nCoV-2019_11_RIGHT | nCoV-2019_1 | TCATCAGATTCAACTTGCATGGCA | 24 | 41.67 | 61.35 |
| nCoV-2019_12_LEFT | nCoV-2019_2 | AAACATGGAGGAGGTGTTGCAG | 22 | 50 | 61.08 |
| nCoV-2019_12_RIGHT | nCoV-2019_2 | TTCACTCTTCATTTCCAAAAAGCTTGA | 27 | 33.33 | 60.36 |
| nCoV-2019_13_LEFT | nCoV-2019_1 | TCGCACAAATGTCTACTTAGCTGT | 24 | 41.67 | 60.56 |

| | | | | | |
|---|---|---|---|---|---|
| nCoV-2019_13_RIGHT | nCoV-2019_1 | ACCACAGCAGTTAAAACACCCT | 22 | 45.45 | 60.36 |
| nCoV-2019_14_LEFT | nCoV-2019_2 | CATCCAGATTCTGCCACTCTTGT | 23 | 47.83 | 60.62 |
| nCoV-2019_14_LEFT_alt4 | nCoV-2019_2 | TGGCAATCTTCATCCAGATTCTGC | 24 | 45.83 | 61.47 |
| nCoV-2019_14_RIGHT | nCoV-2019_2 | AGTTTCCACACAGACAGGCATT | 22 | 45.45 | 60.42 |
| nCoV-2019_14_RIGHT_alt2 | nCoV-2019_2 | TGCGTGTTTCTTCTGCATGTGC | 22 | 50 | 62.76 |
| nCoV-2019_15_LEFT | nCoV-2019_1 | ACAGTGCTTAAAAAGTGTAAAAGTGCC | 27 | 37.04 | 61.32 |
| nCoV-2019_15_LEFT_alt1 | nCoV-2019_1 | AGTGCTTAAAAAGTGTAAAAGTGCCT | 26 | 34.62 | 60.13 |
| nCoV-2019_15_RIGHT | nCoV-2019_1 | AACAGAAACTGTAGCTGGCACT | 22 | 45.45 | 60.16 |
| nCoV-2019_15_RIGHT_alt3 | nCoV-2019_1 | ACTGTAGCTGGCACTTTGAGAGA | 23 | 47.83 | 61.57 |
| nCoV-2019_16_LEFT | nCoV-2019_2 | AATTTGGAAGAAGCTGCTCGGT | 22 | 45.45 | 60.82 |
| nCoV-2019_16_RIGHT | nCoV-2019_2 | CACAACTTGCGTGTGGAGGTTA | 22 | 50 | 61.32 |
| nCoV-2019_17_LEFT | nCoV-2019_1 | CTTCTTTCTTTGAGAGAAGTGAGGACT | 27 | 40.74 | 60.69 |
| nCoV-2019_17_RIGHT | nCoV-2019_1 | TTTGTTGGAGTGTTAACAATGCAGT | 25 | 36 | 60.11 |
| nCoV-2019_18_LEFT | nCoV-2019_2 | TGGAAATACCCACAAGTTAATGGTTTAAC | 29 | 34.48 | 60.69 |
| nCoV-2019_18_LEFT_alt2 | nCoV-2019_2 | ACTTCTATTAAATGGGCAGATAACAACTGT | 30 | 33.33 | 61.38 |
| nCoV-2019_18_RIGHT | nCoV-2019_2 | AGCTTGTTTACCACACGTACAAGG | 24 | 45.83 | 61.51 |
| nCoV-2019_18_RIGHT_alt1 | nCoV-2019_2 | GCTTGTTTACCACACGTACAAGG | 23 | 47.83 | 60.3 |
| nCoV-2019_19_LEFT | nCoV-2019_1 | GCTGTTATGTACATGGGCACACT | 23 | 47.83 | 61.18 |
| nCoV-2019_19_RIGHT | nCoV-2019_1 | TGTCCAACTTAGGGTCAATTTCTGT | 25 | 40 | 60.4 |
| nCoV-2019_20_LEFT | nCoV-2019_2 | ACAAAGAAAACAGTTACACAACAACCA | 27 | 33.33 | 60.68 |
| nCoV-2019_20_RIGHT | nCoV-2019_2 | ACGTGGCTTTATTAGTTGCATTGTT | 25 | 36 | 60.28 |
| nCoV-2019_21_LEFT | nCoV-2019_1 | TGGCTATTGATTATAAACACTACACACCC | 29 | 37.93 | 61.49 |
| nCoV-2019_21_LEFT_alt2 | nCoV-2019_1 | GGCTATTGATTATAAACACTACACACCCT | 29 | 37.93 | 61.29 |
| nCoV-2019_21_RIGHT | nCoV-2019_1 | TAGATCTGTGTGGCCAACCTCT | 22 | 50 | 60.83 |
| nCoV-2019_21_RIGHT_alt0 | nCoV-2019_1 | GATCTGTGTGGCCAACCTCTTC | 22 | 54.55 | 61.2 |
| nCoV-2019_22_LEFT | nCoV-2019_2 | ACTACCGAAGTTGTAGGAGACATTATACT | 29 | 37.93 | 61.25 |
| nCoV-2019_22_RIGHT | nCoV-2019_2 | ACAGTATTCTTTGCTATAGTAGTCGGC | 27 | 40.74 | 60.73 |
| nCoV-2019_23_LEFT | nCoV-2019_1 | ACAACTACTAACATAGTTACACGGTGT | 27 | 37.04 | 60.26 |
| nCoV-2019_23_RIGHT | nCoV-2019_1 | ACCAGTACAGTAGGTTGCAATAGTG | 25 | 44 | 60.57 |
| nCoV-2019_24_LEFT | nCoV-2019_2 | AGGCATGCCTTCTTACTGTACTG | 23 | 47.83 | 60.37 |
| nCoV-2019_24_RIGHT | nCoV-2019_2 | ACATTCTAACCATAGCTGAAATCGGG | 26 | 42.31 | 61.19 |
| nCoV-2019_25_LEFT | nCoV-2019_1 | GCAATTGTTTTTCAGCTATTTTGCAGT | 27 | 33.33 | 60.73 |

| | | | | | |
|---|---|---|---|---|---|
| nCoV-2019_25_RIGHT | nCoV-2019_1 | ACTGTAGTGACAAGTCTCTCGCA | 23 | 47.83 | 61.3 |
| nCoV-2019_26_LEFT | nCoV-2019_2 | TTGTGATACATTCTGTGCTGGTAGT | 25 | 40 | 60.28 |
| nCoV-2019_26_RIGHT | nCoV-2019_2 | TCCGCACTATCACCAACATCAG | 22 | 50 | 60.42 |
| nCoV-2019_27_LEFT | nCoV-2019_1 | ACTACAGTCAGCTTATGTGTCAACC | 25 | 44 | 60.8 |
| nCoV-2019_27_RIGHT | nCoV-2019_1 | AATACAAGCACCAAGGTCACGG | 22 | 50 | 61.13 |
| nCoV-2019_28_LEFT | nCoV-2019_2 | ACATAGAAGTTACTGGCGATAGTTGT | 26 | 38.46 | 60.13 |
| nCoV-2019_28_RIGHT | nCoV-2019_2 | TGTTTAGACATGACATGAACAGGTGT | 26 | 38.46 | 60.91 |
| nCoV-2019_29_LEFT | nCoV-2019_1 | ACTTGTGTTCCTTTTTGTTGCTGC | 24 | 41.67 | 61.39 |
| nCoV-2019_29_RIGHT | nCoV-2019_1 | AGTGTACTCTATAAGTTTTGATGGTGTGT | 29 | 34.48 | 60.69 |
| nCoV-2019_30_LEFT | nCoV-2019_2 | GCACAACTAATGGTGACTTTTTGCA | 25 | 40 | 61.19 |
| nCoV-2019_30_RIGHT | nCoV-2019_2 | ACCACTAGTAGATACACAAACACCAG | 26 | 42.31 | 60.3 |
| nCoV-2019_31_LEFT | nCoV-2019_1 | TTCTGAGTACTGTAGGCACGGC | 22 | 54.55 | 62.03 |
| nCoV-2019_31_RIGHT | nCoV-2019_1 | ACAGAATAAACACCAGGTAAGAATGAGT | 28 | 35.71 | 60.69 |
| nCoV-2019_32_LEFT | nCoV-2019_2 | TGGTGAATACAGTCATGTAGTTGCC | 25 | 44 | 61.09 |
| nCoV-2019_32_RIGHT | nCoV-2019_2 | AGCACATCACTACGCAACTTTAGA | 24 | 41.67 | 60.56 |
| nCoV-2019_33_LEFT | nCoV-2019_1 | ACTTTTGAAGAAGCTGCGCTGT | 22 | 45.45 | 61.58 |
| nCoV-2019_33_RIGHT | nCoV-2019_1 | TGGACAGTAAACTACGTCATCAAGC | 25 | 44 | 61.08 |
| nCoV-2019_34_LEFT | nCoV-2019_2 | TCCCATCTGGTAAAGTTGAGGGT | 23 | 47.83 | 61.02 |
| nCoV-2019_34_RIGHT | nCoV-2019_2 | AGTGAAATTGGGCCTCATAGCA | 22 | 45.45 | 60.03 |
| nCoV-2019_35_LEFT | nCoV-2019_1 | TGTTCGCATTCAACCAGGACAG | 22 | 50 | 61.39 |
| nCoV-2019_35_RIGHT | nCoV-2019_1 | ACTTCATAGCCACAAGGTTAAAGTCA | 26 | 38.46 | 60.69 |
| nCoV-2019_36_LEFT | nCoV-2019_2 | TTAGCTTGGTTGTACGCTGCTG | 22 | 50 | 61.44 |
| nCoV-2019_36_RIGHT | nCoV-2019_2 | GAACAAAGACCATTGAGTACTCTGGA | 26 | 42.31 | 60.74 |
| nCoV-2019_37_LEFT | nCoV-2019_1 | ACACACCACTGGTTGTTACTCAC | 23 | 47.83 | 60.93 |
| nCoV-2019_37_RIGHT | nCoV-2019_1 | GTCCACACTCTCCTAGCACCAT | 22 | 54.55 | 61.48 |
| nCoV-2019_38_LEFT | nCoV-2019_2 | ACTGTGTTATGTATGCATCAGCTGT | 25 | 40 | 60.86 |
| nCoV-2019_38_RIGHT | nCoV-2019_2 | CACCAAGAGTCAGTCTAAAGTAGCG | 25 | 48 | 61.13 |
| nCoV-2019_39_LEFT | nCoV-2019_1 | AGTATTGCCCTATTTTCTTCATAACTGGT | 29 | 34.48 | 61 |
| nCoV-2019_39_RIGHT | nCoV-2019_1 | TGTAACTGGACACATTGAGCCC | 22 | 50 | 60.55 |
| nCoV-2019_40_LEFT | nCoV-2019_2 | TGCACATCAGTAGTCTTACTCTCAGT | 26 | 42.31 | 61.25 |
| nCoV-2019_40_RIGHT | nCoV-2019_2 | CATGGCTGCATCACGGTCAAAT | 22 | 50 | 62.09 |
| nCoV-2019_41_LEFT | nCoV-2019_1 | GTTCCCTTCCATCATATGCAGCT | 23 | 47.83 | 60.75 |

| | | | | | |
|---|---|---|---|---|---|
| nCoV-2019_41_RIGHT | nCoV-2019_1 | TGGTATGACAACCATTAGTTTGGCT | 25 | 40 | 60.75 |
| nCoV-2019_42_LEFT | nCoV-2019_2 | TGCAAGAGATGGTTGTGTTCCC | 22 | 50 | 61.08 |
| nCoV-2019_42_RIGHT | nCoV-2019_2 | CCTACCTCCCTTTGTTGTGTTGT | 23 | 47.83 | 60.69 |
| nCoV-2019_43_LEFT | nCoV-2019_1 | TACGACAGATGTCTTGTGCTGC | 22 | 50 | 60.93 |
| nCoV-2019_43_RIGHT | nCoV-2019_1 | AGCAGCATCTACAGCAAAAGCA | 22 | 45.45 | 61.14 |
| nCoV-2019_44_LEFT | nCoV-2019_2 | TGCCACAGTACGTCTACAAGCT | 22 | 50 | 61.66 |
| nCoV-2019_44_LEFT_alt3 | nCoV-2019_2 | CCACAGTACGTCTACAAGCTGG | 22 | 54.55 | 60.67 |
| nCoV-2019_44_RIGHT | nCoV-2019_2 | AACCTTTCCACATACCGCAGAC | 22 | 50 | 60.87 |
| nCoV-2019_44_RIGHT_alt0 | nCoV-2019_2 | CGCAGACGGTACAGACTGTGTT | 22 | 54.55 | 62.77 |
| nCoV-2019_45_LEFT | nCoV-2019_1 | TACCTACAACTTGTGCTAATGACCC | 25 | 44 | 60.57 |
| nCoV-2019_45_LEFT_alt2 | nCoV-2019_1 | AGTATGTACAAATACCTACAACTTGTGCT | 29 | 34.48 | 60.94 |
| nCoV-2019_45_RIGHT | nCoV-2019_1 | AAATTGTTTCTTCATGTTGGTAGTTAGAGA | 30 | 30 | 60.01 |
| nCoV-2019_45_RIGHT_alt7 | nCoV-2019_1 | TTCATGTTGGTAGTTAGAGAAAGTGTGTC | 29 | 37.93 | 61.53 |
| nCoV-2019_46_LEFT | nCoV-2019_2 | TGTCGCTTCCAAGAAAAGGACG | 22 | 50 | 61.38 |
| nCoV-2019_46_LEFT_alt1 | nCoV-2019_2 | CGCTTCCAAGAAAAGGACGAAGA | 23 | 47.83 | 61.35 |
| nCoV-2019_46_RIGHT | nCoV-2019_2 | CACGTTCACCTAAGTTGGCGTA | 22 | 50 | 60.86 |
| nCoV-2019_46_RIGHT_alt2 | nCoV-2019_2 | CACGTTCACCTAAGTTGGCGTAT | 23 | 47.83 | 61.17 |
| nCoV-2019_47_LEFT | nCoV-2019_1 | AGGACTGGTATGATTTTGTAGAAAACCC | 28 | 39.29 | 61.42 |
| nCoV-2019_47_RIGHT | nCoV-2019_1 | AATAACGGTCAAAGAGTTTTAACCTCTC | 28 | 35.71 | 60.06 |
| nCoV-2019_48_LEFT | nCoV-2019_2 | TGTTGACACTGACTTAACAAAGCCT | 25 | 40 | 61.09 |
| nCoV-2019_48_RIGHT | nCoV-2019_2 | TAGATTACCAGAAGCAGCGTGC | 22 | 50 | 60.74 |
| nCoV-2019_49_LEFT | nCoV-2019_1 | AGGAATTACTTGTGTATGCTGCTGA | 25 | 40 | 60.57 |
| nCoV-2019_49_RIGHT | nCoV-2019_1 | TGACGATGACTTGGTTAGCATTAATACA | 28 | 35.71 | 61.05 |
| nCoV-2019_50_LEFT | nCoV-2019_2 | GTTGATAAGTACTTTGATTGTTACGATGGT | 30 | 33.33 | 60.59 |
| nCoV-2019_50_RIGHT | nCoV-2019_2 | TAACATGTTGTGCCAACCACCA | 22 | 45.45 | 60.95 |
| nCoV-2019_51_LEFT | nCoV-2019_1 | TCAATAGCCGCCACTAGAGGAG | 22 | 54.55 | 61.34 |
| nCoV-2019_51_RIGHT | nCoV-2019_1 | AGTGCATTAACATTGGCCGTGA | 22 | 45.45 | 61.14 |
| nCoV-2019_52_LEFT | nCoV-2019_2 | CATCAGGAGATGCCACAACTGC | 22 | 54.55 | 61.83 |
| nCoV-2019_52_RIGHT | nCoV-2019_2 | GTTGAGAGCAAAATTCATGAGGTCC | 25 | 44 | 60.62 |
| nCoV-2019_53_LEFT | nCoV-2019_1 | AGCAAAATGTTGGACTGAGACTGA | 24 | 41.67 | 60.69 |
| nCoV-2019_53_RIGHT | nCoV-2019_1 | AGCCTCATAAAACTCAGGTTCCC | 23 | 47.83 | 60.31 |
| nCoV-2019_54_LEFT | nCoV-2019_2 | TGAGTTAACAGGACACATGTTAGACA | 26 | 38.46 | 60.18 |

| | | | | | |
|---|---|---|---|---|---|
| nCoV-2019_54_RIGHT | nCoV-2019_2 | AACCAAAAACTTGTCCATTAGCACA | 25 | 36 | 60.11 |
| nCoV-2019_55_LEFT | nCoV-2019_1 | ACTCAACTTTACTTAGGAGGTATGAGCT | 28 | 39.29 | 61.43 |
| nCoV-2019_55_RIGHT | nCoV-2019_1 | GGTGTACTCTCCTATTTGTACTTTACTGT | 29 | 37.93 | 60.54 |
| nCoV-2019_56_LEFT | nCoV-2019_2 | ACCTAGACCACCACTTAACCGA | 22 | 50 | 60.49 |
| nCoV-2019_56_RIGHT | nCoV-2019_2 | ACACTATGCGAGCAGAAGGGTA | 22 | 50 | 61.21 |
| nCoV-2019_57_LEFT | nCoV-2019_1 | ATTCTACACTCCAGGGACCACC | 22 | 54.55 | 61.16 |
| nCoV-2019_57_RIGHT | nCoV-2019_1 | GTAATTGAGCAGGGTCGCCAAT | 22 | 50 | 61.26 |
| nCoV-2019_58_LEFT | nCoV-2019_2 | TGATTTGAGTGTTGTCAATGCCAGA | 25 | 40 | 61.44 |
| nCoV-2019_58_RIGHT | nCoV-2019_2 | CTTTTCTCCAAGCAGGGTTACGT | 23 | 47.83 | 61.06 |
| nCoV-2019_59_LEFT | nCoV-2019_1 | TCACGCATGATGTTTCATCTGCA | 23 | 43.48 | 61.42 |
| nCoV-2019_59_RIGHT | nCoV-2019_1 | AAGAGTCCTGTTACATTTTCAGCTTG | 26 | 38.46 | 60.02 |
| nCoV-2019_60_LEFT | nCoV-2019_2 | TGATAGAGACCTTTATGACAAGTTGCA | 27 | 37.04 | 60.53 |
| nCoV-2019_60_RIGHT | nCoV-2019_2 | GGTACCAACAGCTTCTCTAGTAGC | 24 | 50 | 60.44 |
| nCoV-2019_61_LEFT | nCoV-2019_1 | TGTTTATCACCCGCGAAGAAGC | 22 | 50 | 61.5 |
| nCoV-2019_61_RIGHT | nCoV-2019_1 | ATCACATAGACAACAGGTGCGC | 22 | 50 | 61.25 |
| nCoV-2019_62_LEFT | nCoV-2019_2 | GGCACATGGCTTTGAGTTGACA | 22 | 50 | 61.91 |
| nCoV-2019_62_RIGHT | nCoV-2019_2 | GTTGAACCTTTCTACAAGCCGC | 22 | 50 | 60.35 |
| nCoV-2019_63_LEFT | nCoV-2019_1 | TGTTAAGCGTGTTGACTGGACT | 22 | 45.45 | 60.16 |
| nCoV-2019_63_RIGHT | nCoV-2019_1 | ACAAACTGCCACCATCACAACC | 22 | 50 | 61.85 |
| nCoV-2019_64_LEFT | nCoV-2019_2 | TCGATAGATATCCTGCTAATTCCATTGT | 28 | 35.71 | 60.11 |
| nCoV-2019_64_RIGHT | nCoV-2019_2 | AGTCTTGTAAAAGTGTTCCAGAGGT | 25 | 40 | 60.1 |
| nCoV-2019_65_LEFT | nCoV-2019_1 | GCTGGCTTTAGCTTGTGGGTTT | 22 | 50 | 61.92 |
| nCoV-2019_65_RIGHT | nCoV-2019_1 | TGTCAGTCATAGAACAAACACCAATAGT | 28 | 35.71 | 60.9 |
| nCoV-2019_66_LEFT | nCoV-2019_2 | GGGTGTGGACATTGCTGCTAAT | 22 | 50 | 61.21 |
| nCoV-2019_66_RIGHT | nCoV-2019_2 | TCAATTTCCATTTGACTCCTGGGT | 24 | 41.67 | 60.45 |
| nCoV-2019_67_LEFT | nCoV-2019_1 | GTTGTCCAACAATTACCTGAAACTTACT | 28 | 35.71 | 60.43 |
| nCoV-2019_67_RIGHT | nCoV-2019_1 | CAACCTTAGAAACTACAGATAAATCTTGGG | 30 | 36.67 | 60.4 |
| nCoV-2019_68_LEFT | nCoV-2019_2 | ACAGGTTCATCTAAGTGTGTGTGT | 24 | 41.67 | 60.14 |
| nCoV-2019_68_RIGHT | nCoV-2019_2 | CTCCTTTATCAGAACCAGCACCA | 23 | 47.83 | 60.31 |
| nCoV-2019_69_LEFT | nCoV-2019_1 | TGTCGCAAAATATACTCAACTGTGTCA | 27 | 37.04 | 61.43 |
| nCoV-2019_69_RIGHT | nCoV-2019_1 | TCTTTATAGCCACGGAACCTCCA | 23 | 47.83 | 61.14 |
| nCoV-2019_70_LEFT | nCoV-2019_2 | ACAAAAGAAAATGACTCTAAAGAGGGTTT | 29 | 31.03 | 60.13 |

| | | | | | |
|---|---|---|---|---|---|
| nCoV-2019_70_RIGHT | nCoV-2019_2 | TGACCTTCTTTTAAAGACATAACAGCAG | 28 | 35.71 | 60.27 |
| nCoV-2019_71_LEFT | nCoV-2019_1 | ACAAATCCAATTCAGTTGTCTTCCTATTC | 29 | 34.48 | 60.54 |
| nCoV-2019_71_RIGHT | nCoV-2019_1 | TGGAAAAGAAAGGTAAGAACAAGTCCT | 27 | 37.04 | 60.8 |
| nCoV-2019_72_LEFT | nCoV-2019_2 | ACACGTGGTGTTTATTACCCTGAC | 24 | 45.83 | 61.04 |
| nCoV-2019_72_RIGHT | nCoV-2019_2 | ACTCTGAACTCACTTTCCATCCAAC | 25 | 44 | 60.97 |
| nCoV-2019_73_LEFT | nCoV-2019_1 | CAATTTTGTAATGATCCATTTTTGGGTGT | 29 | 31.03 | 60.29 |
| nCoV-2019_73_RIGHT | nCoV-2019_1 | CACCAGCTGTCCAACCTGAAGA | 22 | 54.55 | 62.45 |
| nCoV-2019_74_LEFT | nCoV-2019_2 | ACATCACTAGGTTTCAAACTTTACTTGC | 28 | 35.71 | 60.68 |
| nCoV-2019_74_RIGHT | nCoV-2019_2 | GCAACACAGTTGCTGATTCTCTTC | 24 | 45.83 | 60.85 |
| nCoV-2019_75_LEFT | nCoV-2019_1 | AGAGTCCAACCAACAGAATCTATTGT | 26 | 38.46 | 60.24 |
| nCoV-2019_75_RIGHT | nCoV-2019_1 | ACCACCAACCTTAGAATCAAGATTGT | 26 | 38.46 | 60.69 |
| nCoV-2019_76_LEFT | nCoV-2019_2 | AGGGCAAACTGGAAAGATTGCT | 22 | 45.45 | 60.76 |
| nCoV-2019_76_LEFT_alt3 | nCoV-2019_2 | GGGCAAACTGGAAAGATTGCTGA | 23 | 47.83 | 61.87 |
| nCoV-2019_76_RIGHT | nCoV-2019_2 | ACACCTGTGCCTGTTAAACCAT | 22 | 45.45 | 60.42 |
| nCoV-2019_76_RIGHT_alt0 | nCoV-2019_2 | ACCTGTGCCTGTTAAACCATTGA | 23 | 43.48 | 60.69 |
| nCoV-2019_77_LEFT | nCoV-2019_1 | CCAGCAACTGTTTGTGGACCTA | 22 | 50 | 60.75 |
| nCoV-2019_77_RIGHT | nCoV-2019_1 | CAGCCCCTATTAAACAGCCTGC | 22 | 54.55 | 61.59 |
| nCoV-2019_78_LEFT | nCoV-2019_2 | CAACTTACTCCTACTTGGCGTGT | 23 | 47.83 | 60.55 |
| nCoV-2019_78_RIGHT | nCoV-2019_2 | TGTGTACAAAAACTGCCATATTGCA | 25 | 36 | 60.22 |
| nCoV-2019_79_LEFT | nCoV-2019_1 | GTGGTGATTCAACTGAATGCAGC | 23 | 47.83 | 60.92 |
| nCoV-2019_79_RIGHT | nCoV-2019_1 | CATTTCATCTGTGAGCAAAGGTGG | 24 | 45.83 | 60.62 |
| nCoV-2019_80_LEFT | nCoV-2019_2 | TTGCCTTGGTGATATTGCTGCT | 22 | 45.45 | 60.89 |
| nCoV-2019_80_RIGHT | nCoV-2019_2 | TGGAGCTAAGTTGTTTAACAAGCG | 24 | 41.67 | 60.02 |
| nCoV-2019_81_LEFT | nCoV-2019_1 | GCACTTGGAAAACTTCAAGATGTGG | 25 | 44 | 61.24 |
| nCoV-2019_81_RIGHT | nCoV-2019_1 | GTGAAGTTCTTTTCTTGTGCAGGG | 24 | 45.83 | 60.73 |
| nCoV-2019_82_LEFT | nCoV-2019_2 | GGGCTATCATCTTATGTCCTTCCCT | 25 | 48 | 61.52 |
| nCoV-2019_82_RIGHT | nCoV-2019_2 | TGCCAGAGATGTCACCTAAATCAA | 24 | 41.67 | 60.02 |
| nCoV-2019_83_LEFT | nCoV-2019_1 | TCCTTTGCAACCTGAATTAGACTCA | 25 | 40 | 60.46 |
| nCoV-2019_83_RIGHT | nCoV-2019_1 | TTTGACTCCTTTGAGCACTGGC | 22 | 50 | 61.33 |
| nCoV-2019_84_LEFT | nCoV-2019_2 | TGCTGTAGTTGTCTCAAGGGCT | 22 | 50 | 61.61 |
| nCoV-2019_84_RIGHT | nCoV-2019_2 | AGGTGTGAGTAAACTGTTACAAACAAC | 27 | 37.04 | 60.36 |
| nCoV-2019_85_LEFT | nCoV-2019_1 | ACTAGCACTCTCCAAGGGTGTT | 22 | 50 | 61.03 |

| | | | | | |
|---|---|---|---|---|---|
| nCoV-2019_85_RIGHT | nCoV-2019_1 | ACACAGTCTTTTACTCCAGATTCCC | 25 | 44 | 60.51 |
| nCoV-2019_86_LEFT | nCoV-2019_2 | TCAGGTGATGGCACAACAAGTC | 22 | 50 | 61.07 |
| nCoV-2019_86_RIGHT | nCoV-2019_2 | ACGAAAGCAAGAAAAAGAAGTACGC | 25 | 40 | 61.01 |
| nCoV-2019_87_LEFT | nCoV-2019_1 | CGACTACTAGCGTGCCTTTGTA | 22 | 50 | 60.16 |
| nCoV-2019_87_RIGHT | nCoV-2019_1 | ACTAGGTTCCATTGTTCAAGGAGC | 24 | 45.83 | 60.81 |
| nCoV-2019_88_LEFT | nCoV-2019_2 | CCATGGCAGATTCCAACGGTAC | 22 | 54.55 | 61.58 |
| nCoV-2019_88_RIGHT | nCoV-2019_2 | TGGTCAGAATAGTGCCATGGAGT | 23 | 47.83 | 61.4 |
| nCoV-2019_89_LEFT | nCoV-2019_1 | GTACGCGTTCCATGTGGTCATT | 22 | 50 | 61.5 |
| nCoV-2019_89_LEFT_alt2 | nCoV-2019_1 | CGCGTTCCATGTGGTCATTCAA | 22 | 50 | 62.01 |
| nCoV-2019_89_RIGHT | nCoV-2019_1 | ACCTGAAAGTCAACGAGATGAAACA | 25 | 40 | 60.91 |
| nCoV-2019_89_RIGHT_alt4 | nCoV-2019_1 | ACGAGATGAAACATCTGTTGTCACT | 25 | 40 | 60.74 |
| nCoV-2019_90_LEFT | nCoV-2019_2 | ACACAGACCATTCCAGTAGCAGT | 23 | 47.83 | 61.58 |
| nCoV-2019_90_RIGHT | nCoV-2019_2 | TGAAATGGTGAATTGCCCTCGT | 22 | 45.45 | 60.82 |
| nCoV-2019_91_LEFT | nCoV-2019_1 | TCACTACCAAGAGTGTGTTAGAGGT | 25 | 44 | 60.93 |
| nCoV-2019_91_RIGHT | nCoV-2019_1 | TTCAAGTGAGAACCAAAAGATAATAAGCA | 29 | 31.03 | 60.03 |
| nCoV-2019_92_LEFT | nCoV-2019_2 | TTTGTGCTTTTTAGCCTTTCTGCT | 24 | 37.5 | 60.14 |
| nCoV-2019_92_RIGHT | nCoV-2019_2 | AGGTTCCTGGCAATTAATTGTAAAAGG | 27 | 37.04 | 60.53 |
| nCoV-2019_93_LEFT | nCoV-2019_1 | TGAGGCTGGTTCTAAATCACCCA | 23 | 47.83 | 61.59 |
| nCoV-2019_93_RIGHT | nCoV-2019_1 | AGGTCTTCCTTGCCATGTTGAG | 22 | 50 | 60.55 |
| nCoV-2019_94_LEFT | nCoV-2019_2 | GGCCCCAAGGTTTACCCAATAA | 22 | 50 | 60.56 |
| nCoV-2019_94_RIGHT | nCoV-2019_2 | TTTGGCAATGTTGTTCCTTGAGG | 23 | 43.48 | 60.18 |
| nCoV-2019_95_LEFT | nCoV-2019_1 | TGAGGGAGCCTTGAATACACCA | 22 | 50 | 61.1 |
| nCoV-2019_95_RIGHT | nCoV-2019_1 | CAGTACGTTTTTGCCGAGGCTT | 22 | 50 | 61.95 |
| nCoV-2019_96_LEFT | nCoV-2019_2 | GCCAACAACAACAAGGCCAAAC | 22 | 50 | 61.82 |
| nCoV-2019_96_RIGHT | nCoV-2019_2 | TAGGCTCTGTTGGTGGGAATGT | 22 | 50 | 61.36 |
| nCoV-2019_97_LEFT | nCoV-2019_1 | TGGATGACAAAGATCCAAATTTCAAAGA | 28 | 32.14 | 60.22 |
| nCoV-2019_97_RIGHT | nCoV-2019_1 | ACACACTGATTAAAGATTGCTATGTGAG | 28 | 35.71 | 60.17 |
| nCoV-2019_98_LEFT | nCoV-2019_2 | AACAATTGCAACAATCCATGAGCA | 24 | 37.5 | 60.5 |
| nCoV-2019_98_RIGHT | nCoV-2019_2 | TTCTCCTAAGAAGCTATTAAAATCACATGG | 30 | 33.33 | 60.01 |

**Supplemental Table 4**. Household transmission pair metadata including accession numbers, difference in days between symptom onset, difference in days between collection dates, and pair identifier.

| C | Column 3 | Column 4 | Column 5 | Column 6 | Column 7 | Column 8 | Column 9 | Column 10 | Column 11 | Column 12 | Column 13 | Column 14 |
|---|---|---|---|---|---|---|---|---|---|---|---|---|
| 0 | tip1 | tip2 | muts_between_tips | muts | probability_1_serial_inteval | tube_IDs | time_between_test | direction_based_on_test_date | time_between_symptoms | direction_based_on_symptoms | #comparisons | pair_number |
| 0 | USA/WI-UW-41/2020 | USA/WI-UW-48/2020 | 0 | [] | 0.7496528051576963 | 20,28 | 0 | 28 <--> 20 | 1 | 28 <--> 20 | 2 | pair1, pair1a (28 --> 20), pair 1b (20 --> 28) |
| 0 | USA/WI-UW-65/2020 | USA/WI-UW-32/2020 | 0 | [] | 0.7496528051576963 | 50,8 | 2 | 8 <--> 50 | 2 | 8 <--> 50 | 2 | pair2, pair2a (8 --> 50), pair2b (50 --> 8) |
| 0 | USA/WI-UW-69/2020 | USA/WI-UW-61/2020 | 0 | [] | 0.7496528051576963 | 55,44 | 4 | 55 --> 44 | 3 | 55 <--> 44 | 2 | pair3, pair3a (55 --> 44), pair3b (44 --> 55) |
| 0 | USA/WI-UW-70/2020 | USA/WI-UW-67/2020 | 0 | [] | 0.7496528051576963 | 56,53 | 6 | 56 --> 53 | 5 | 56 --> 53 | 1 | pair4 |
| 0 | USA/WI-UW-74/2020 | USA/WI-UW-29/2020 | 0 | [] | 0.7496528051576963 | 61,5 | 4 | 61 --> 5 | 4 | 61 --> 5 | 1 | pair5 |
| 0 | USA/WI-UW-438/202 | USA/WI-UW-432/202 | 0 | [] | 0.7496528051576963 | 744,738 | 2 | 738 --> 744 | 2 | 738 --> 744 | 2 | pair6, pair 6a (738 --> 744), pair 6b (744 --> 738) |
| 0 | USA/WI-UW-544/202 | USA/WI-UW-551/202 | 1 | ['T4917C'] | 0.21600878707411836 | 893,884 | 1 | 893 <--> 884 | 4 | 884 --> 893 | 1 | pair7 |
| 0 | USA/WI-UW-544/202 | USA/WI-UW-575/202 | 0 | [] | 0.7496528051576963 | 884,903 | 1 | 884 <--> 903 | asx | 884 <--> 903 | 2 | pair8, pair8a (884 --> 903), pair8b (903 --> 884) |
| 0 | USA/WI-UW-551/202 | USA/WI-UW-575/202 | 1 | ['T4917C'] | 0.21600878707411836 | 893,903 | 0 | 893 <--> 903 | asx | 893 <--> 903 | 2 | pair9, pair9a (893 --> 903), pair9b (903 --> 893) |
| 0 | USA/WI-UW-546/202 | USA/WI-UW-586/202 | 0 | [] | 0.7496528051576963 | 887,916 | 0 | 887 <--> 916 | 1 | 887 <--> 916 | 2 | pair10, pair10a (887 --> 916), pair 10b (916 --> 887) |
| 0 | USA/WI-UW-546/202 | USA/WI-UW-443/202 | 0 | [] | 0.7496528051576963 | 887,749 | 0 | 887 <--> 749 | 0 | 887 <--> 749 | 2 | pair11, pair 11a (887 --> 749), pair 11b (749 --> 887) |
| 0 | USA/WI-UW-586/202 | USA/WI-UW-443/202 | 0 | [] | 0.7496528051576963 | 916,749 | 0 | 916 <--> 749 | 1 | 916 <--> 749 | 2 | pair12, pair12a (916 --> 749), pair12b (749 --> 916) |
| 0 | USA/WI-UW-577/202 | USA/WI-UW-536/202 | 0 | [] | 0.7496528051576963 | 906,849 | 0 | 906 <--> 849 | asx | 906 <--> 849 | 2 | pair13, pair13a (906 --> 849), piar13b (849 --> 906) |
| 0 | USA/WI-UW-598/202 | USA/WI-UW-602/202 | 0 | [] | 0.7496528051576963 | 956,962 | 3 | 956 --> 962 | 4 | 962 --> 956 | 2 | pair14 |
| 0 | USA/WI-UW-601/202 | USA/WI-UW-780/202 | 0 | [] | 0.7496528051576963 | 961,1195 | 8 | 961 --> 1195 | 5 | 961 --> 1195 | 1 | pair15 |
| ?( | USA/WI-UW-756/202 | USA/WI-UW-893/202 | 0 | [] | 0.7496528051576963 | 1157,1346 | 7 | 1157 --> 1346 | 6 | 1157 --> 1346 | 1 | pair16 |
| 0 | USA/WI-UW-874/202 | USA/WI-UW-986/202 | 0 | [] | 0.7496528051576963 | 1326,1495 | 3 | 1326 --> 1495 | asx | 1326 <--> 1495 | 1 | pair17 |
| 0 | USA/WI-UW-874/202 | USA/WI-UW-997/202 | 0 | [] | 0.7496528051576963 | 1326,1512 | 3 | 1326 --> 1512 | asx | 1326 <--> 1512 | 1 | pair18 |
| 0 | USA/WI-UW-874/202 | USA/WI-UW-991/202 | 0 | [] | 0.7496528051576963 | 1326,1502 | 3 | 1326 --> 1502 | asx | 1326 <--> 1502 | 1 | pair19 |
| 0 | USA/WI-UW-986/202 | USA/WI-UW-997/202 | 0 | [] | 0.7496528051576963 | 1495,1512 | 0 | 1495 <--> 1512 | asx | 1495 <--> 1512 | 2 | pair20, pair20a (1495 --> 1512), pair20b (1512 --> 1495) |
| 0 | USA/WI-UW-986/202 | USA/WI-UW-991/202 | 0 | [] | 0.7496528051576963 | 1495,1502 | 0 | 1495 <--> 1502 | asx | 1495 <--> 1502 | 2 | pair21, pair21a (1495 --> 1502), pair21b (1502 --> 1495) |
| 0 | USA/WI-UW-997/202 | USA/WI-UW-991/202 | 0 | [] | 0.7496528051576963 | 1512,1502 | 0 | 1512 <--> 1502 | asx | 1512 <--> 1502 | 2 | pair22, pair22a (1512 --> 1502), pair22b (1502 --> 1512) |
| 0 | USA/WI-UW-895/202 | USA/WI-UW-876/202 | 0 | [] | 0.7496528051576963 | 1353,1328 | 0 | 1353 <--> 1328 | 3 | 1353 --> 1328 | 1 | pair23 |
| 0 | USA/WI-UW-895/202 | USA/WI-UW-863/202 | 2 | ['A15942C', 'C25006T'] | 0.031120937434107567 | 1353,1297 | 0 | 1353 <--> 1297 | 3 | 1353 --> 1297 | 1 | pair24 |
| 0 | USA/WI-UW-876/202 | USA/WI-UW-863/202 | 2 | ['A15942C', 'C25006T'] | 0.031120937434107567 | 1328,1297 | 0 | 1297 <--> 1328 | 0 | 1297 <--> 1328 | 1 | pair25, pair25a (1297 --> 1328), pair25b (1328 --> 1297) |
| 0 | USA/WI-UW-158/202 | USA/WI-UW-160/202 | 0 | [] | 0.7496528051576963 | 195,197 | 0 | 195 <--> 197 | NA | 195 <--> 197 | 2 | pair26, pair26a (195 --> 197), pair26b (197 --> 195) |
| 0 | USA/WI-UW-333/202 | USA/WI-UW-334/202 | 0 | [] | 0.7496528051576963 | 453,454 | 0 | 453 <--> 454 | NA | 453 <--> 454 | 2 | pair27, pair27a (453 --> 454), pair27b (454 --> 453) |
| 0 | USA/WI-UW-119/202 | USA/WI-UW-120/202 | 0 | [] | 0.7496528051576963 | 128,130 | 3 | 128 --> 130 | 10 | 130 --> 128 | 1 | pair28 |
